# A criterion characterizing accumulated neurotoxicity of Aβ oligomers in Alzheimer’s disease

**DOI:** 10.1101/2024.08.19.608707

**Authors:** Andrey V. Kuznetsov

## Abstract

The paper develops a criterion to quantify the accumulated neurotoxicity of Aβ oligomers in Alzheimer’s disease (AD). Accumulated neurotoxicity is determined by integrating the concentration of Aβ oligomers within the control volume over time. In the scenario of a low rate of free Aβ oligomer deposition into senile plaques and dysfunctional degradation machinery, resulting in an infinitely long half-life of Aβ monomers and aggregates, the obtained analytical solution reveals a quadratic relationship between accumulated neurotoxicity and time. This suggests that initially, neurotoxicity increases slowly but accelerates as time progresses. This could help to understand the prolonged delay in the onset of AD symptoms. Furthermore, as the model indicates that accumulated neurotoxicity increases with the duration of the aggregation process, it implies that if the protein degradation system is compromised, the onset of AD becomes unavoidable. Eventually, neuronal death is only a question of time. The only way to prevent this outcome is to ensure that the degradation machinery for Aβ peptides and their aggregates remains functional. A threshold value of accumulated neurotoxicity is suggested. The developed theory suggests that if this value is exceeded, nearby neurons will die. The progression of accumulated neurotoxicity over time is analyzed. An S-shaped growth pattern as the half-deposition time of Aβ aggregates into senile plaques increases is revealed. Additionally, the sensitivity of accumulated neurotoxicity to different parameter values is examined.

## 1. Introduction

Alzheimer’s disease (AD) is a progressive neurodegenerative disorder and the leading cause of dementia, characterized by extracellular amyloid beta (Aβ) plaques and intracellular tau tangles. The disease significantly impacts daily life and independence [1-5].

AD involves the aggregation of Aβ monomers into fibrillar structures through secondary nucleation, an autocatalytic process [6]. Aβ monomers are generated by the cleavage of amyloid precursor protein (APP) by β- and γ-secretases, a process that occurs on lipid membranes [7,8]. Most Aβ monomers are then released into the extracellular space, where they can aggregate [9]. In AD Aβ monomers aggregate to form oligomers, which eventually deposit into Aβ plaques.

The “neurotoxic (hereafter toxic) Aβ oligomer” hypothesis suggests that small, soluble oligomers of amyloid peptides cause cellular toxicity, providing a potential explanation for the weak correlation between amyloid plaques and neurodegeneration in AD. In this hypothesis, Aβ plaques are considered relatively inert sinks of toxic intermediate species [10]. Aβ monomers also exhibit minimal toxic activity [11]. The primary hypothesis for AD toxicity is that toxic Aβ oligomers disrupt the neuronal membrane [12]. However, the absence of a quantifiable parameter characterizing oligomer toxicity complicates the determination of a toxic threshold that must be exceeded for neuronal death processes to begin [10].

Mathematical models exploring the mechanisms behind Aβ plaque formation were developed in refs. [13-16]. The Finke-Watzky (F-W) two-step model [17] was employed to simulate Aβ aggregation. Building on this foundation, the present paper introduces a model for calculating the accumulated toxicity of Aβ oligomers. Both numerical and analytical solutions are presented to clarify how accumulated toxicity depends on disease duration and various biophysical processes represented by the model parameters.

## 2. Materials and models

### 2.1 Equations of the mathematical model

The F-W model represents a simplified two-step mechanism for aggregate formation, comprising nucleation and autocatalysis. In the nucleation step, new aggregates form continuously, while in the autocatalysis step, existing aggregates facilitate their own further production [17,18]. The model outlines these two pseudo-elementary reaction steps as follows:

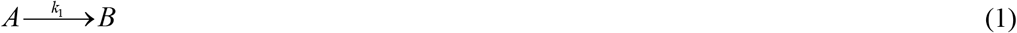

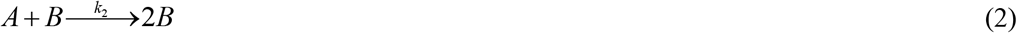

Here, *A* represents a monomeric peptide, while *B* represents a free (not yet deposited into a plaque) amyloid-converted peptide. The kinetic constants *k*_1_ and *k*_2_ denote the rates of nucleation and autocatalytic growth, respectively [17]. Primary nucleation, described by Eq. (1), involves only monomers, whereas secondary nucleation, described by Eq. (2), includes both monomers and existing free aggregates of the same peptide [6].

If Aβ monomers diffuse rapidly, the concentration of Aβ monomers, *C*_*A*_, is nearly uniform in the control volume (CV) shown in Fig. 1. In this scenario, time, *t*, becomes the sole independent variable in the model. The dependent variables utilized in the model are detailed in Table 1, while the parameters are outlined in Table 2.

**Table 1.** Dependent variables utilized in the model.

| Symbol | Definition | Units |
| --- | --- | --- |
| $C_A$ | Concentration of Aβ monomers | $\mu\text{M}$ |
| $C_B$ | Concentration of free Aβ aggregates | $\mu\text{M}$ |
| $C_D$ | Concentration of Aβ aggregates deposited into Aβ plaques | $\mu\text{M}$ |

**Table 2.**
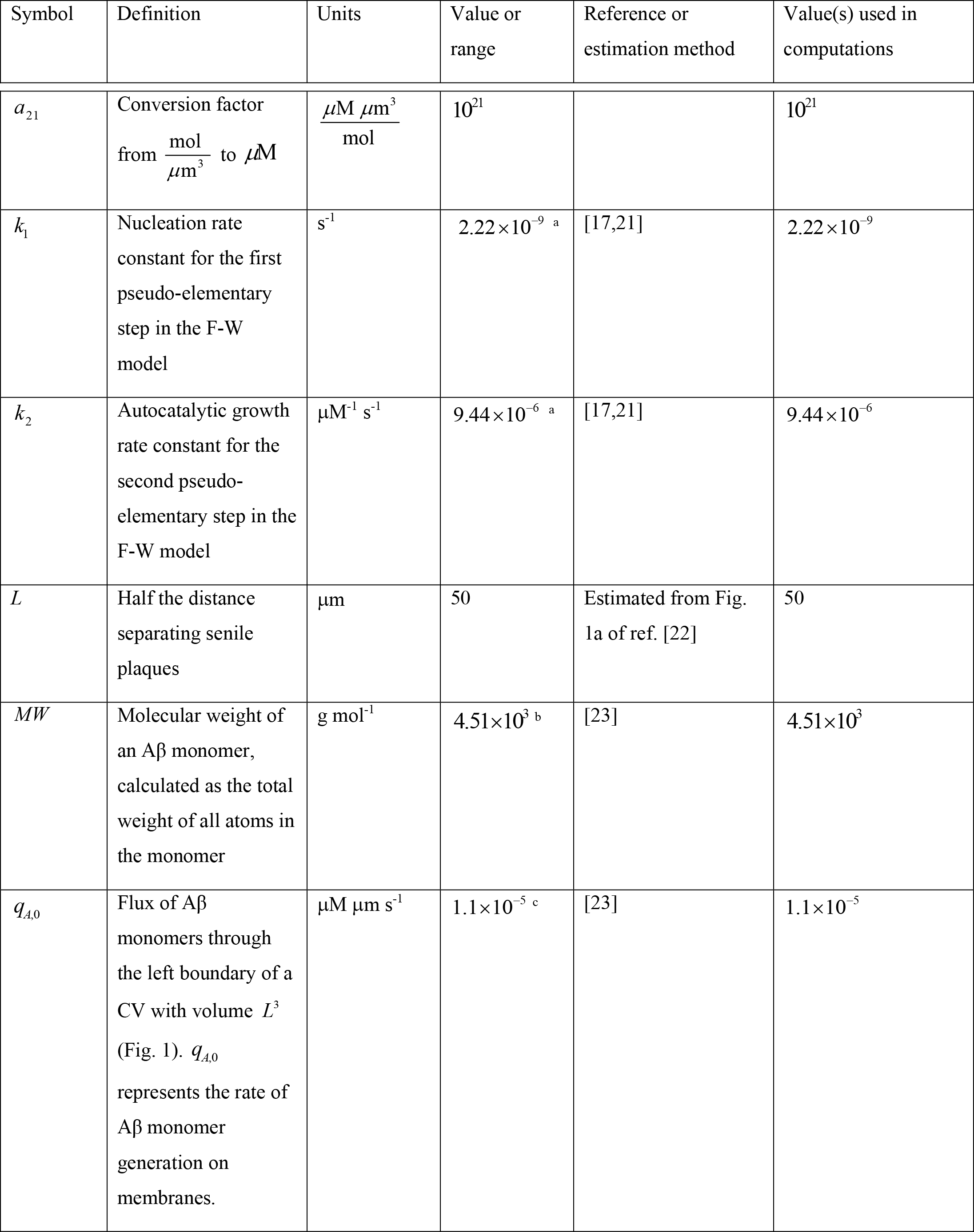
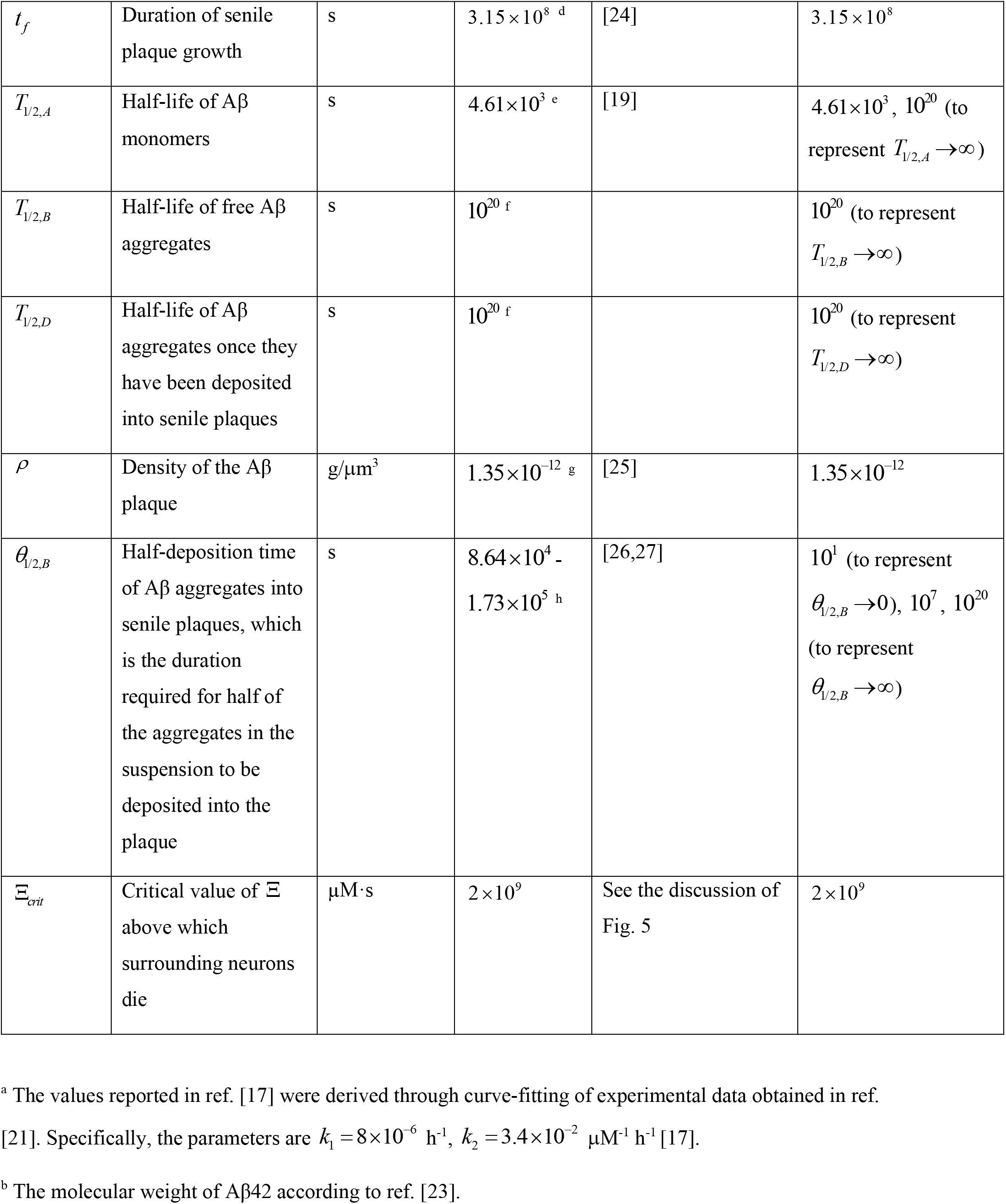
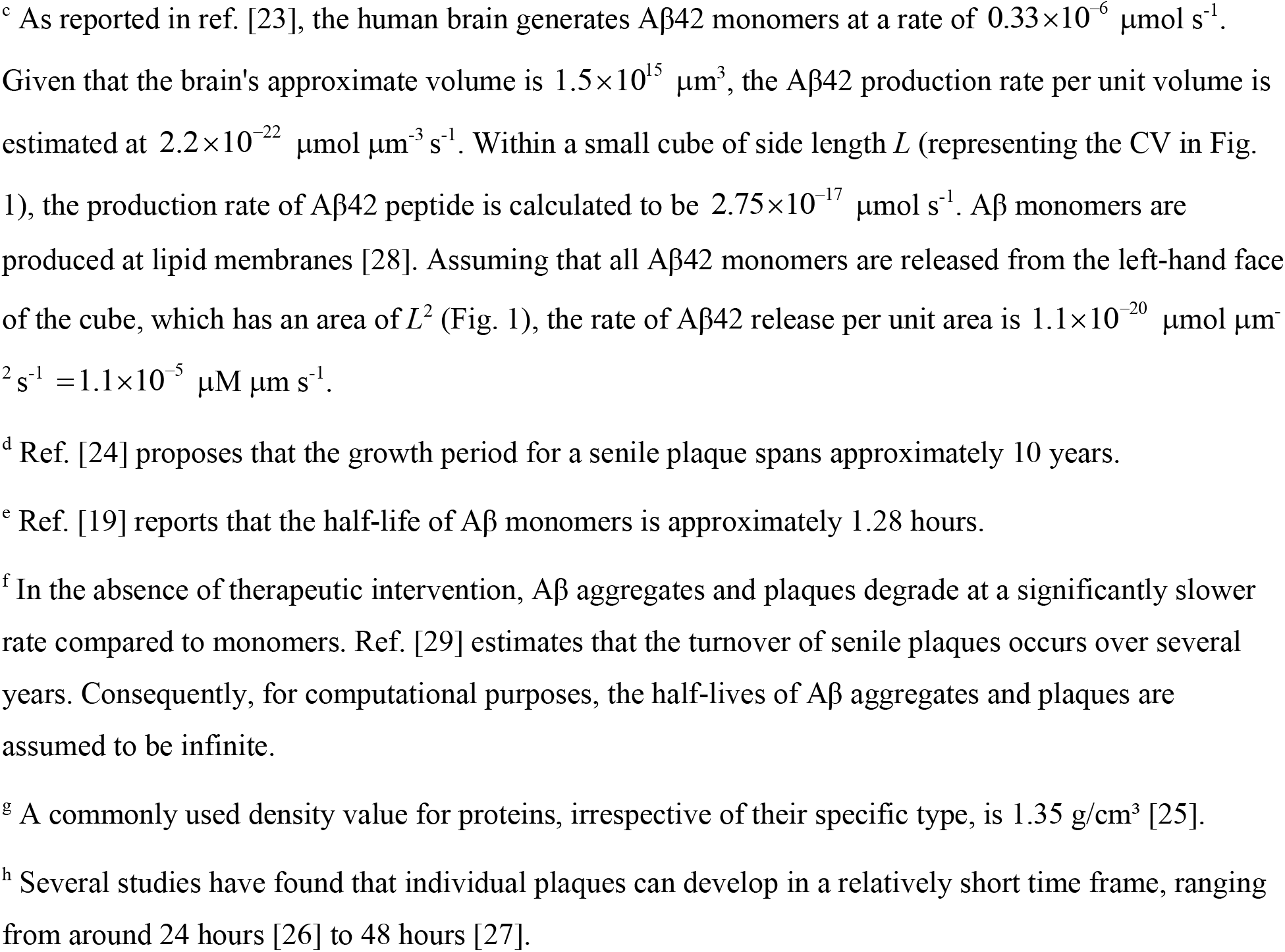
Model parameters.

**Fig. 1.**
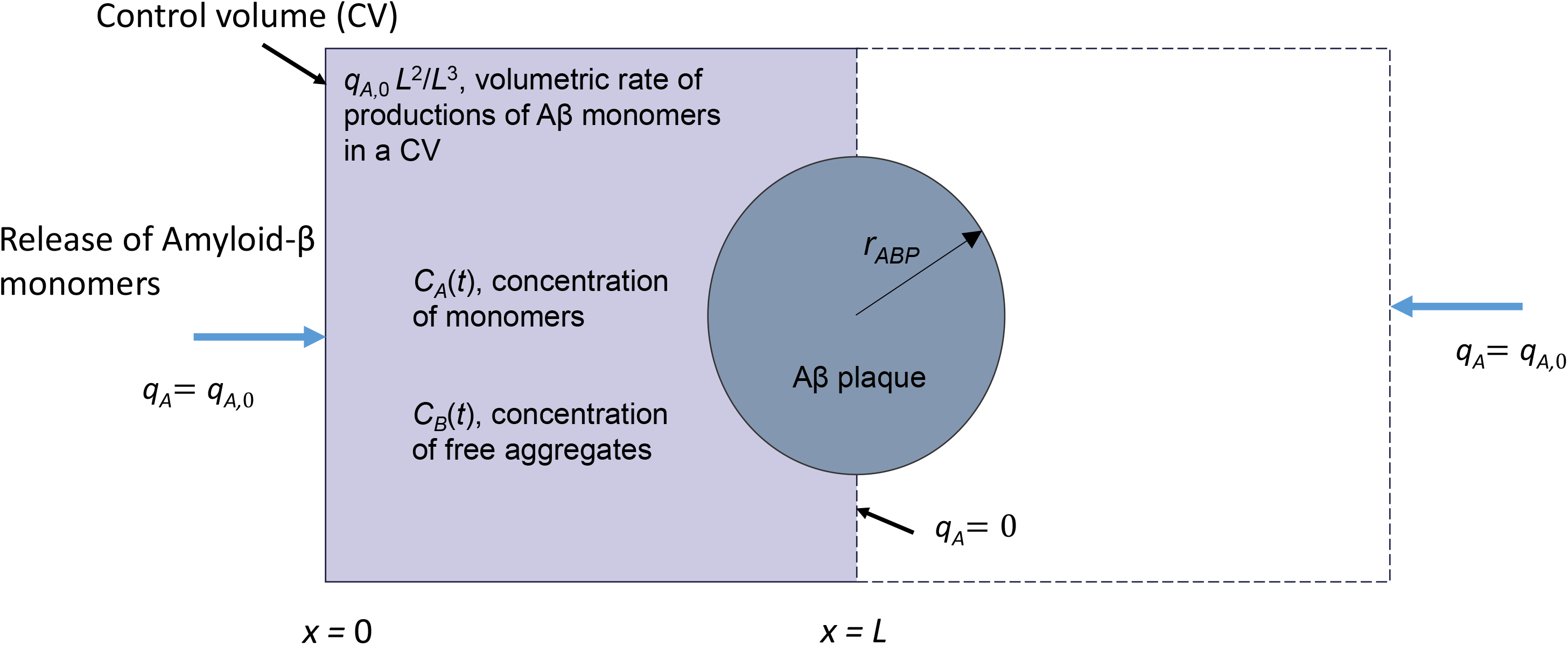
A control volume (CV) with side length *L* used to simulate the aggregation of Aβ monomers into free aggregates and their subsequent deposition into senile plaques. Aβ monomers are assumed to be produced at the *x*=0 boundary. The boundary at *x*=*L* is treated as symmetric, preventing any flux of Aβ monomers. The formation of an Aβ plaque is assumed to occur at the symmetric interface between two adjacent CVs. A lumped capacitance approximation is employed, allowing the assumption that all Aβ concentrations are functions of time only.

By scaling model equations, ref. [15] established the following dimensionless parameter:

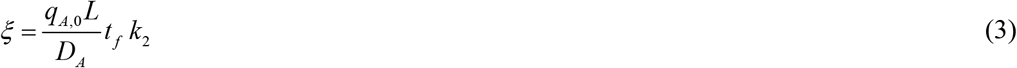

Parameter *ξ* represents the ratio of the variation in Aβ monomer concentration across the CV to the average concentration of Aβ monomers within the CV at time *t*_*f*_, which represents the duration of the process. Ref. [15] demonstrated that the lumped capacitance approximation, which assumes that Aβ concentrations depend only on time and not on location within the CV, holds when *ξ* ≪ 1. In Eq. (3), *D*_*A*_ denotes the diffusivity of Aβ monomers. Using a value of 62.3 μm^2^/s for *D*_*A*_ [19] and taking other parameters from Table 2, *ξ* is estimated to be 0.026. Therefore, the criterion *ξ* ≪ 1 is met, allowing variations in Aβ monomer concentrations across the CV to be neglected.

By applying the conservation of Aβ monomers within the CV shown in Fig. 1, the following equation is derived:

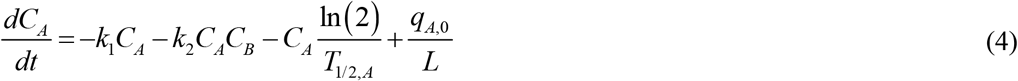

The first term on the right-hand side of Eq. (4) represents the nucleation-driven conversion rate of Aβ monomers into aggregates, while the second term describes the rate of conversion through autocatalytic growth. Additionally, the third term on the right-hand side of Eq. (4) accounts for the degradation of Aβ monomers, and the fourth term simulates the production of Aβ monomers on lipid membranes.

By stating the conservation of free Aβ aggregates within the CV, the following equation is obtained:

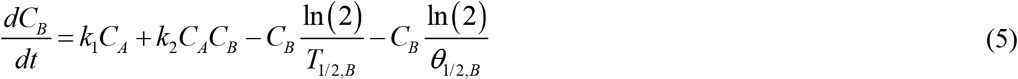

In Eq. (5), the first two terms on the right-hand side are analogous to the first two terms in Eq. (4), but with opposite signs. The third term represents the degradation rate of free Aβ aggregates, while the fourth term models the deposition rate of these aggregates into senile plaques.

The formation of Aβ plaques from adhesive Aβ fibrils is modeled by a method similar to that used for colloidal suspension coagulation [20]. In this model, free Aβ aggregates (*B*) are assumed to deposit into Aβ plaques, and the concentration of these deposited aggregates is denoted as *C*_*D*_.

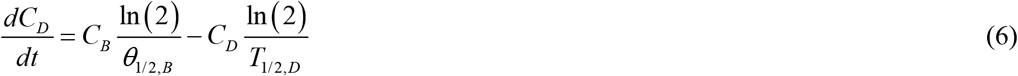

In Eq. (6), the first term on the right-hand side corresponds to the fourth term in Eq. (5) but with the opposite sign. The second term reflects the half-life of Aβ aggregates within the senile plaques.

The initial conditions are defined as follows:

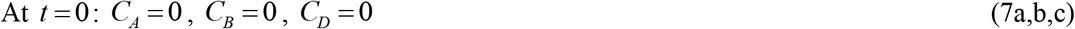

### 2.2. Criterion of accumulated toxicity of Aβ oligomers

The exact mechanism behind the toxicity of Aβ oligomers is not well understood. Certain Aβ oligomers may be more toxic than others. Alternatively, a mixture of different oligomers with varying structures, stability, and concentrations might cause nonspecific toxicity by binding to membrane proteins, targeting membrane lipids, inducing oxidative stress, and altering membrane dielectric properties and ion permeability [10]. If an “Aβ soup” is responsible for nonspecific toxicity, the parameter characterizing the accumulated toxicity of Aβ is

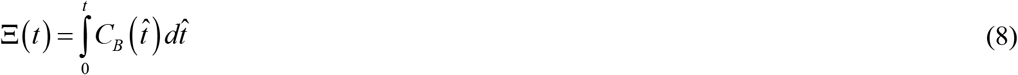

The dimensionless version of Eqs. (4)-(8) is given in section S1.1 of the Supplementary Materials.

If toxicity is caused by several specific toxic oligomer species, the criterion for toxicity can be redefined as follows (for example):

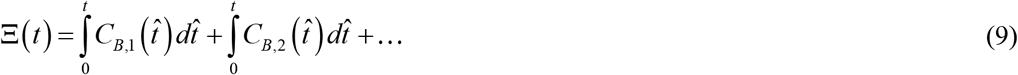

### 2.3. A model for senile plaque growth

A methodology developed in ref. [14] is followed. To simulate the growth of an Aβ plaque (illustrated in Fig. 1), the approach involves determining the total number of Aβ monomers, represented by *N*, that have been incorporated into the plaque over time *t*. This can be done by using the average concentration of Aβ monomers deposited into the plaque, as suggested in ref. [30]:

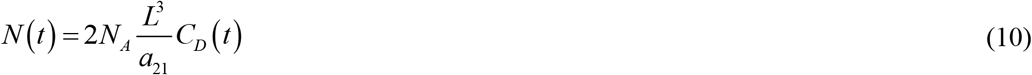

where *N*_*A*_ is Avogadro’s number. On the right-hand side of Eq. (10) the coefficient of 2 arises from the model’s assumption that the Aβ plaque forms between two adjacent neurons. Each neuron’s membrane releases Aβ monomers at a rate of *q*_*A*,0_, contributing to the plaque’s development (Fig. 1).

Another approach, which also follows ref. [30], is determining *N* (*t*) through the volume of a single inclusion body, *V*_*ABP*_ :

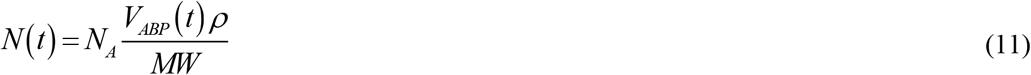

where *MW* denotes the average molecular weight of an Aβ monomer.

By equating the right-hand sides of Eqs. (10) and (11) and solving for the volume of the Aβ plaque, the following expression is derived:

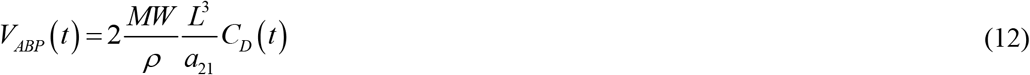

where 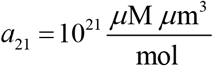 is the conversion factor from 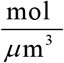 to *μ*M.

Assuming the Aβ plaque takes on a spherical shape,

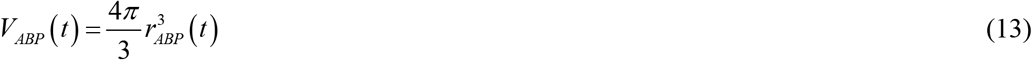

By equating the right sides of Eqs. (12) and (13) and solving for the radius of the Aβ plaque, the following expression is derived:

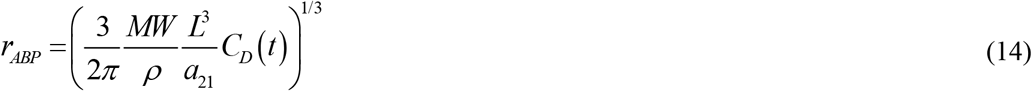

## 3. Analytical solutions for the limiting cases

### 3.1. The scenario of a slow deposition rate of Aβ aggregates into senile plaques, *θ*_1/2,*B*_ →∞, **as well as** *T*_1/2, *A*_ →∞ **and** *T*_1/2, *B*_ →∞

For a slow deposition rate of Aβ aggregates into senile plaques and infinitely large half-lives of monomers and free aggregates, Eqs. (4)-(6) reduce to, respectively,

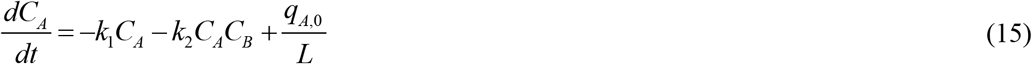

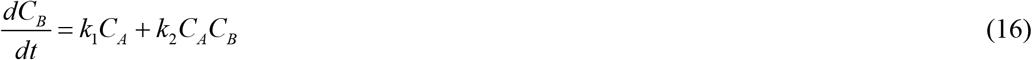

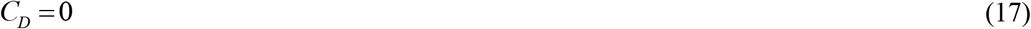

Eqs. (15) and (16) were solved using the initial conditions (7a,b), as detailed in refs. [14,31]. For completeness, the analytical solutions to Eqs. (15) and (16) are provided in section S1.2 of the Supplementary Materials.

The exact solution given by Eqs. (S10) and (S12) is rather cumbersome. A more elegant approximate solution is

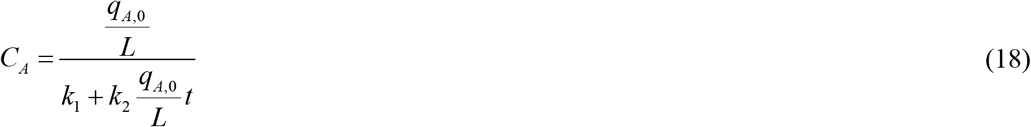

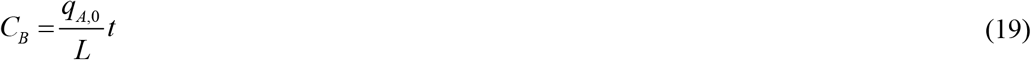

Note that as *t* →∞, Eq. (18) leads to *C*_*A*_ → 0.

Using Eqs. (8) and (19),

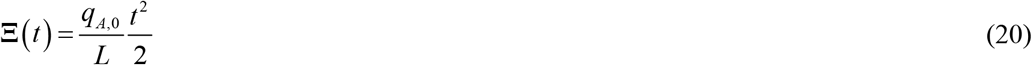

The quadratic dependence on time indicates that initially the accumulated toxicity grows slowly, but as time progresses, it increases more rapidly.

### 3.2. The scenario of a fast deposition rate of Aβ aggregates into senile plaques, *θ*_1/2,*B*_ → 0, **as well as** *T*_1/ 2, *A*_ →∞ **and** *T*_1/ 2, *B*_ →∞

As *θ*_1/2,*B*_ → 0, which means immediate deposition of Aβ aggregates into plaques, Eqs. (4)-(6) simplify to, respectively,

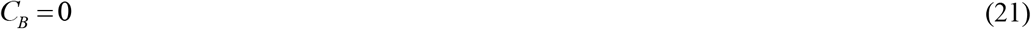

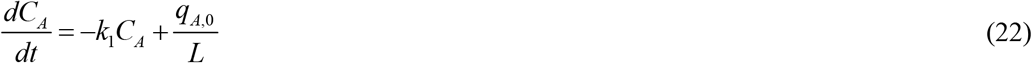

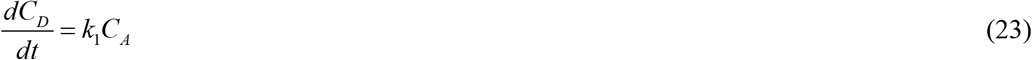

In this case, Eqs. (8) and (21) lead to

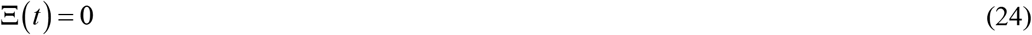

Adding Eqs. (22) and (23) gives

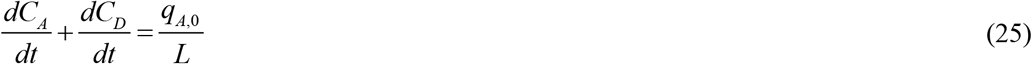

Integrating Eq. (25) over time and applying the initial conditions from Eq. (7a,c), the following result is derived:

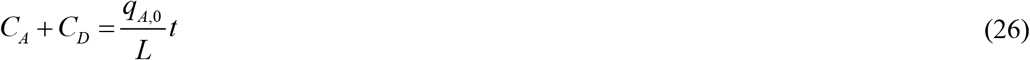

The increase in *C*_*A*_ + *C*_*D*_ over time is due to the influx of newly synthesized Aβ monomers at the left (or right) boundary of the CV, as shown in Fig. 1. By substituting *C*_*A*_ from Eq. (26) into Eq. (23), the following is derived:

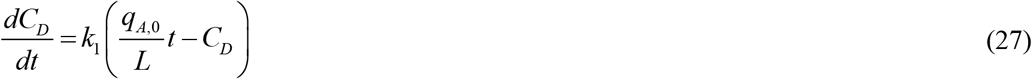

The exact solution of Eq. (27) with the initial condition specified in Eq. (7c) is

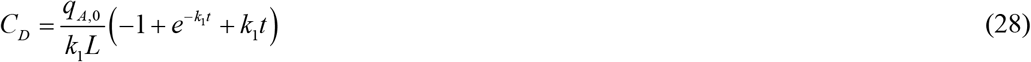

As *t* → 0, Eq. (28) becomes

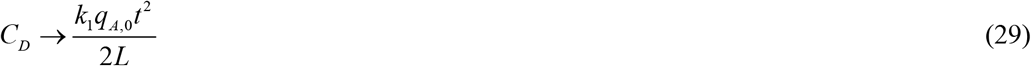

indicating that at small *t*, the concentration of Aβ aggregates deposited into Aβ plaques is directly proportional to the kinetic constant characterizing the nucleation of Aβ aggregates.

The quantity *C*_*A*_ can then be determined using Eqs. (26) and (28) as

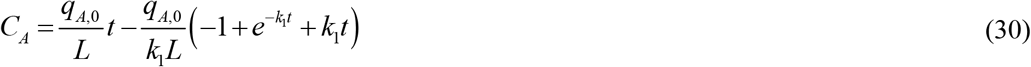

## 4. Sensitivity analysis

One of the primary objectives of this study was to examine the sensitivity of accumulated toxicity, Ξ, in response to various parameters.

This involved calculating the local sensitivity coefficients, which are the first-order partial derivatives of the observable (the accumulated toxicity) with respect to parameters such as the nucleation rate constant for the first pseudo-elementary step in the F-W model, *k*_1_. This analysis followed the methodology outlined in refs. [32-35]. Specifically, the sensitivity coefficient of Ξ to a parameter (for example) *k*_1_ was determined using

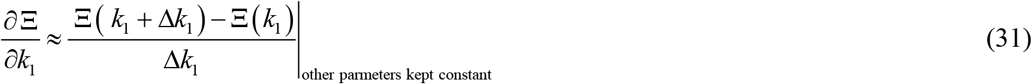

where Δ*k*_1_ = 10^−3^ *k*_1_ is the step size. Calculations were conducted with various step sizes to evaluate the sensitivity coefficients’ independence from the step size.

The non-dimensional relative sensitivity coefficients were computed following the method described in refs. [33,36]. For example, the coefficient characterizing the sensitivity of accumulated toxicity Ξ to the nucleation rate constant *k*_1_ is defined as:

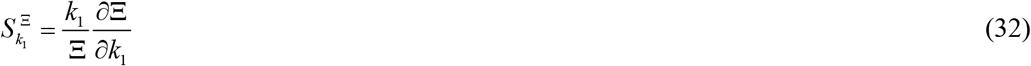

From Eq. (20), the following is obtained:

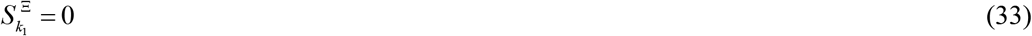

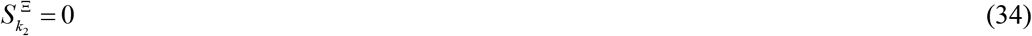

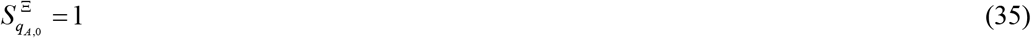

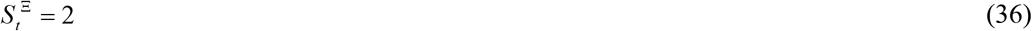

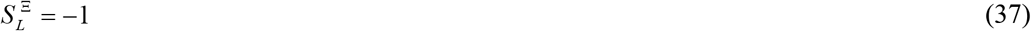

## 5. Results

Details of the numerical solution methodology are provided in Section S1.3 of the Supplementary Materials.

Figs. S1 and S2 display the concentrations of Aβ monomers, *C*_*A*_ (*t*), free Aβ aggregates, *C*_*B*_ (*t*), Aβ aggregates deposited in plaques, *C*_*D*_ (*t*), and the radius of a growing Aβ plaque, *r*_*ABP*_ (*t*), for different values of *k*_1_. Interestingly, *C*_*A*_ appears to be independent of *k*_1_ (Fig. S1a). Indeed, in this scenario, the last two terms in Eq. (4) are dominant. By equating these terms, the following estimate of *C*_*A*_ is derived:

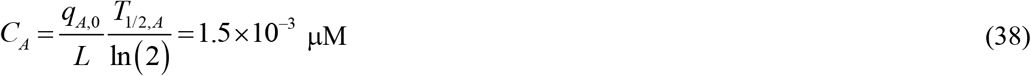

This corresponds to the value that *C*_*A*_ rapidly reaches in Fig. S1a. Note that the value derived from Eq. (38) is independent of *k*_1_.

The value that *C*_*B*_ reaches in Fig. S1b depends on *k*_1_ because the production rate of free aggregates through nucleation, *k*_1_*C*_*A*_, is directly proportional to *k*_1_ (refer to Eq. (5)).

Since free aggregates deposit into plaques (see Eq. (6)), an increase in the nucleation rate constant *k*_1_ also leads to a rise in the concentration of deposited aggregates, *C*_*D*_ (Fig. S2a). Notably, *C*_*D*_ increases linearly over time (Fig. S2a). This can be attributed to the assumed constant rate of Aβ monomer production, *q*_*A*,0_, and the fact that the rate of Aβ monomer degradation, 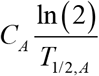, remains unaffected by *k* in this scenario.

Since the radius of the plaque is proportional to the cube root of the concentration of deposited aggregates (see Eq. (14)), *r*_*ABP*_ increases in proportion to the cube root of time, *t*_^1/3^_. The increase in *k*_1_ also leads to an increase in the Aβ plaque radius *r*_*ABP*_ (Fig. S2b).

Figs. S3 and S4 display concentrations *C*_*A*_ (*t*), *C*_*B*_ (*t*), *C*_*D*_ (*t*), and *r*_*ABP*_ (*t*) for various values of the autocatalytic growth rate constant *k*_2_. The dependence of *C*_*A*_ (*t*) on *t* becomes more complex; for the greatest value of *k*_2_ the monomer concentration decreases as *t* increases (Fig. S3a). This occurs because *k*_2_ controls the production of free Aβ aggregates from monomers via autocatalytic growth, represented by the nonlinear term, *k*_2_*C*_*A*_*C*_*B*_, in Eq. (4). This nonlinear process cannot be easily balanced by the linear term governing Aβ monomer degradation, 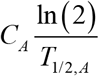, in Eq. (4). The values of *C*_*B*_, *C*_*D*_, and *r*_*ABP*_ increase as *k*_2_ increases, but now in a nonlinear manner, increasing much faster for larger values of *k*_2_ (Figs. S3b, S4a, and S4b).

Figs. S5 and S6 display *C*_*A*_ (*t*), *C*_*B*_ (*t*), *C*_*D*_ (*t*), and *r*_*ABP*_ (*t*) for different values of the production rate of Aβ monomers, *q*_*A*,0_. The behavior of these quantities with respect to *q*_*A*,0_ is somewhat similar to its dependence on *k*_2_ ; *C*_*A*_ (*t*) decreases with *t* for large values of *q*_*A*,0_ (Fig. S5a). Meanwhile, *C*_*B*_, *C*_*D*_, and *r*_*ABP*_ increase as *q*_*A*,0_ increases, with this growth accelerating significantly for greater values of *q*_*A*,0_.

In Fig. 2a, the numerically computed accumulated toxicity, Ξ, increases linearly over time. For *θ* _1/2,*B*_ = 10^7^ s, the increase in *k*_1_ leads to a corresponding increase in Ξ (Fig. 2a). The increase remains slow when *k*_1_ is small but accelerates as *k*_1_ increases. As *θ*_1/2,*B*_ →∞, Ξ becomes independent of *k*_1_ and aligns with the analytical solution given by Eq. (20) (Fig. 2b). It is important to note that the analytical solution predicts that Ξ ∼ *t*^2^.

**Fig. 2.**
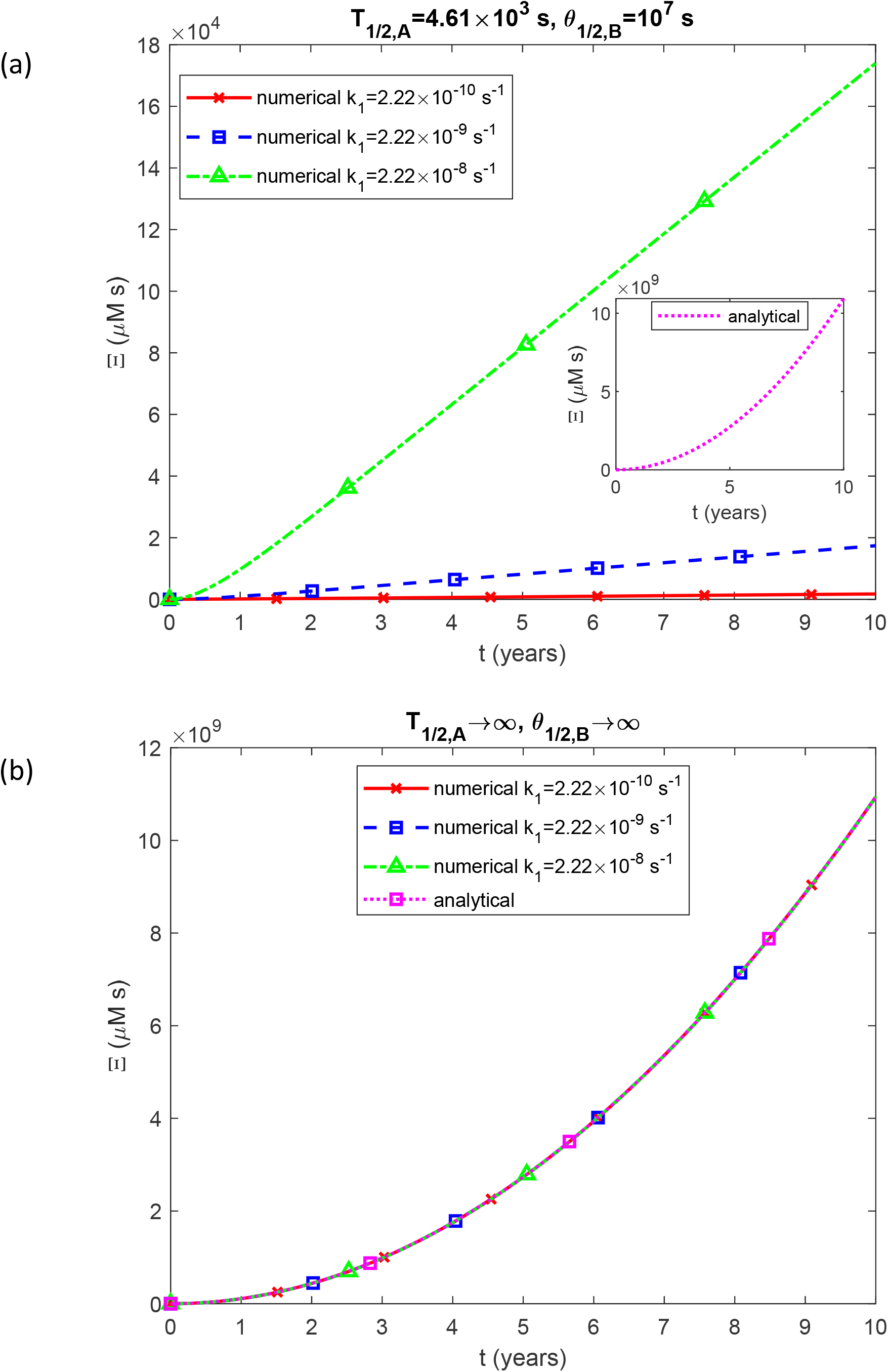
Accumulated toxicity of Aβ oligomers, Ξ, as a function of time (in years). Results are shown for three different values of *k*_1_. (a) *T*_1/ 2, *A*_ = 4.61×10^3^ and *θ* _1/2,*B*_ =10^7^ s, (b) *T*_1/ 2, *A*_ →∞ and *θ* _1/2,*B*_ →∞. Note that as *θ*_1/2,*B*_ → 0, all free aggregates immediately deposit into plaques, resulting in Ξ→0. Note that the analytical data do not depend on *k*_1_.

At low values of *k*_2_, the accumulated toxicity, Ξ, remains small but increases rapidly when *k*_2_ becomes large (Fig. 3a). In the case where *θ*_1/2,*B*_ →∞, Ξ becomes independent of *k*_2_ and coincides with the analytical result from Eq. (20) (Fig. 3b).

**Fig. 3.**
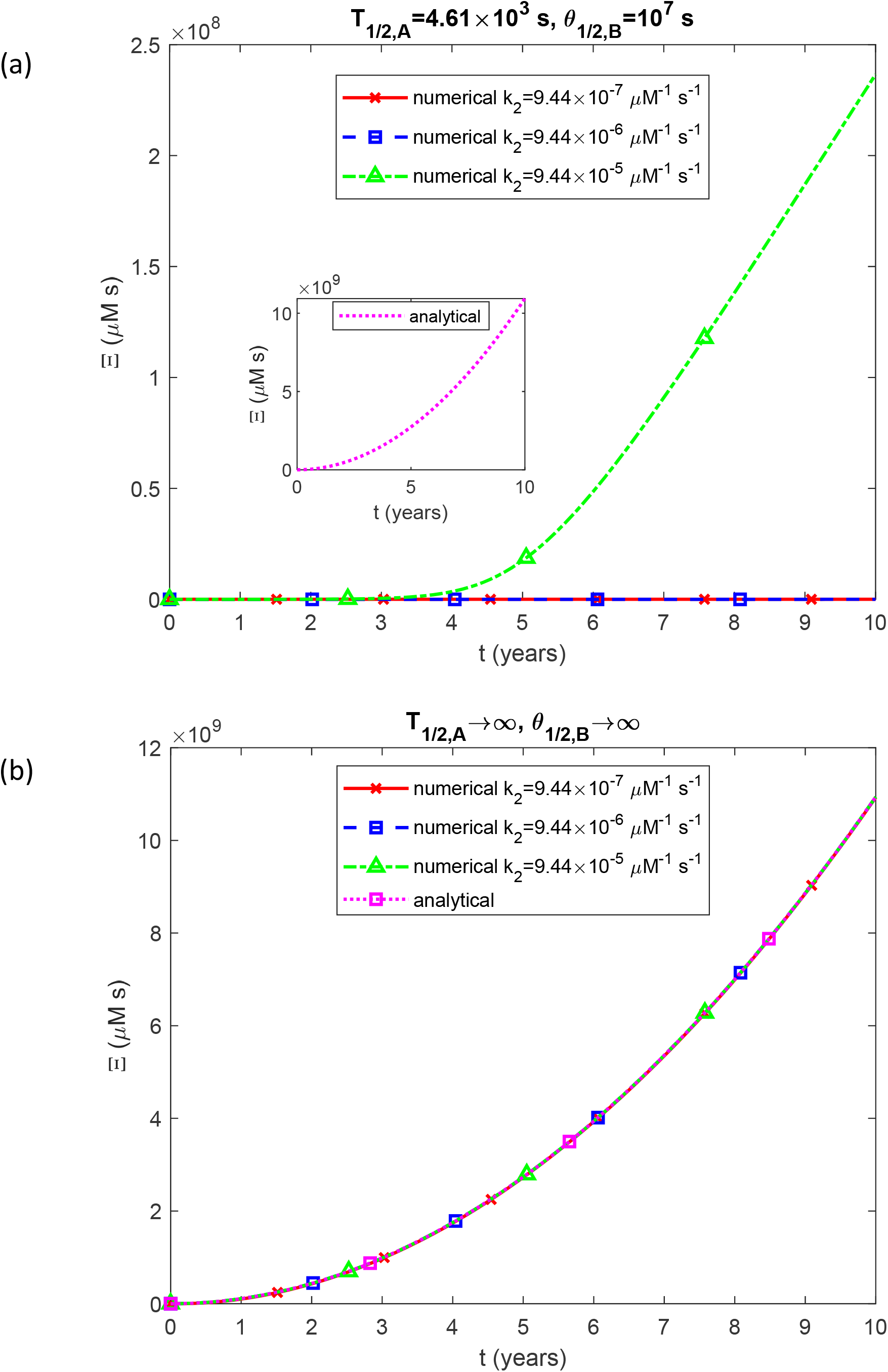
Accumulated toxicity of Aβ oligomers, Ξ, as a function of time (in years). Results are shown for three different values of *k*_2_. (a) *T*_1/ 2, *A*_ = 4.61×10^3^ s and *θ* _1/2,*B*_ =10^7^ s, (b) *T*_1/ 2, *A*_ →∞ and *θ*_1/2,*B*_ →∞. Note that as *θ*_1/2,*B*_ → 0, all free aggregates immediately deposit into plaques, resulting in Ξ→0. Note that the analytical data do not depend on *k*_2_.

When *q*_*A*,0_ is small, the accumulated toxicity, Ξ, stays low but surges dramatically as *q*_*A*,0_ increases to its maximum value in Fig. 4a. As *θ*_1/2,*B*_ →∞, Ξ aligns with the analytical prediction from Eq. (20). In this scenario, Ξ exhibits a linear dependence on *q*_*A*,0_ (see Eq. (20)), explaining the different lines observed for different values of *q*_*A*,0_ in Fig. 4b.

**Fig. 4.**
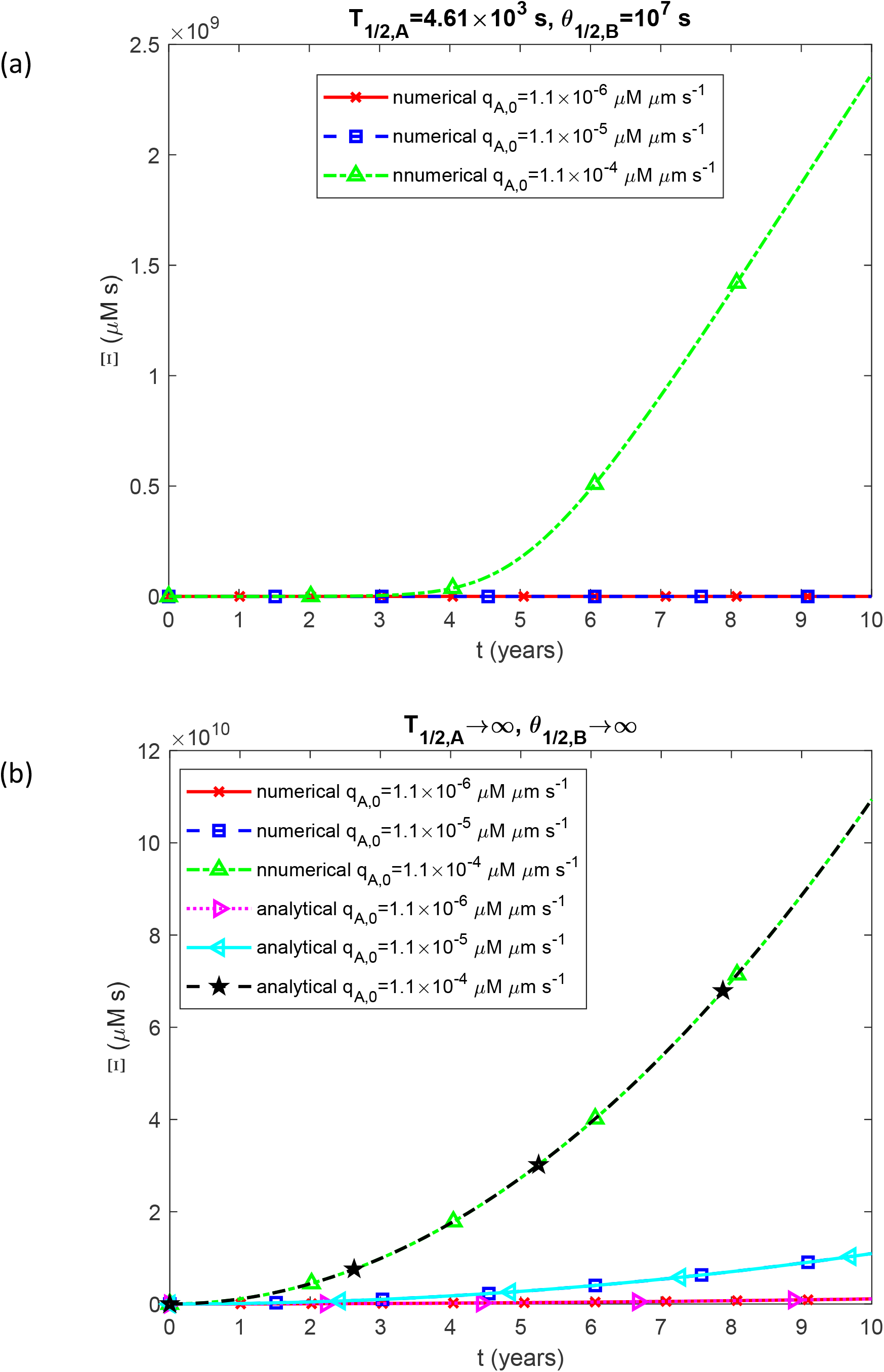
Accumulated toxicity of Aβ oligomers, Ξ, as a function of time (in years). Results are shown for three different values of *q*_*A*,0_. (a) *T*_1/ 2, *A*_ = 4.61×10^3^ s and *θ* _1/2,*B*_ =10^7^ s, (b) *T*_1/ 2, *A*_ →∞ and *θ*_1/2,*B*_ →∞. Note that as *θ*_1/2,*B*_ → 0, all free aggregates immediately deposit into plaques, resulting in Ξ→0.

The results shown in Fig. 4a, illustrating Ξ(*t*), and Fig. S6b, depicting *r*_*ABP*_ (*t*), have been combined to create Fig. 5, which presents the plot of accumulated toxicity vs Aβ plaque radius, Ξ= *f* (*r*_*ABP*_). Despite recent criticisms of the amyloid hypothesis in Alzheimer’s disease (AD) by some authors [37,38], many researchers still believe there is a correlation between the accumulation of sticky amyloid proteins in the brain and AD [39]. Assuming the largest plaque sizes correspond to the most advanced stages of the disease and that nearby neurons die when plaques reach a significant size, the threshold value for Ξ can be estimated from Fig. 5 at approximately 2 ×10^9^ µM·s, which corresponds to the plaque radius of approximately 3 µm.

**Fig. 5.**
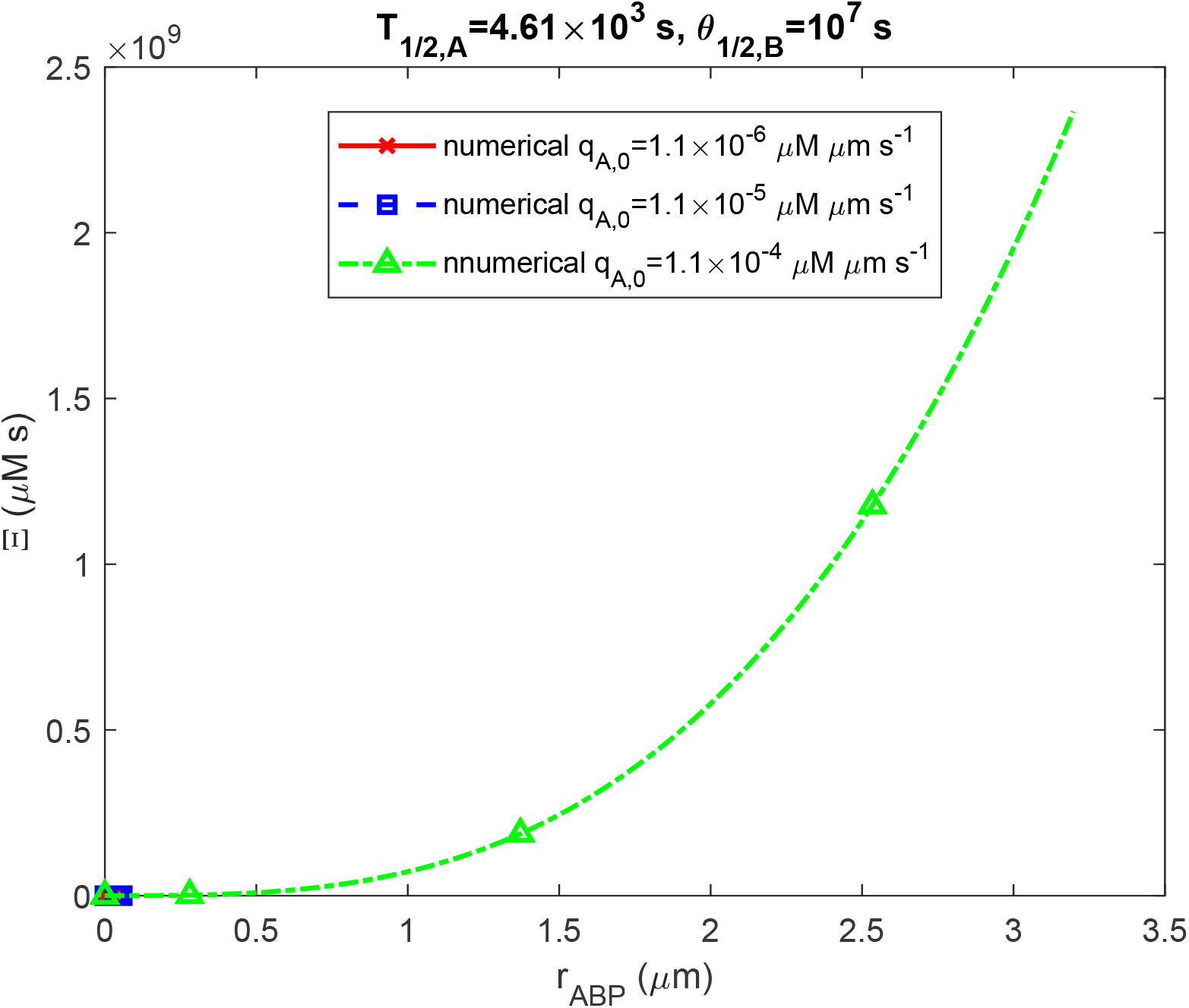
Accumulated toxicity of Aβ oligomers, Ξ, as a function of the radius of a growing Aβ plaque, *r*_*ABP*_. Results are shown for three different values of *q*_*A*,0_. The depicted scenario assumes a finite half-life of Aβ monomers (*T*_1/ 2, *A*_ = 4.61×10^3^ s) and an intermediate value of half-deposition time for Aβ aggregates into senile plaques (*θ*_1/2,*B*_ =10^7^ s). Note that the curves corresponding to smaller values of *q*_*A*,0_ stop at lower values of *r*_*ABP*_ and thus end shortly after the origin.

As the half-deposition time of Aβ aggregates into senile plaques *θ*_1/2,*B*_ →0, free aggregates immediately deposit into plaques, leading to Ξ → 0. The question arises as to what happens with Ξ when *θ*_1/2,*B*_ is increased. This issue is explored in Figs. 6-8. For a finite half-life of Aβ monomers, *T*_1/ 2, *A*_, the accumulated toxicity exhibits a rapid S-shaped increase around *θ*_1/2,*B*_ ≈10^7^ s and then it plateaus, reaching a level still lower than the analytical prediction in Eq. (20) (Fig. 6a). However, as *T*_1/2,*A*_ →∞, Ξ reaches the value predicted by the analytical solution at higher *θ*_1/2,*B*_ values (Fig. 6b). For finite *T*_1/ 2, *A*_, the numerical curves differ for various *k*_1_, but as *T*_1/2,*A*_ →∞, the curves converge (compare Fig. 6a and 6b). Notably, some curves in Fig. 6 cross the solid black horizontal line, which represents the critical value of Ξ, 2 ×10^9^ µM·s. This indicates that in the simulated CV, the accumulated toxicity exceeds its critical value, resulting in the death of nearby neurons.

**Fig. 6.**
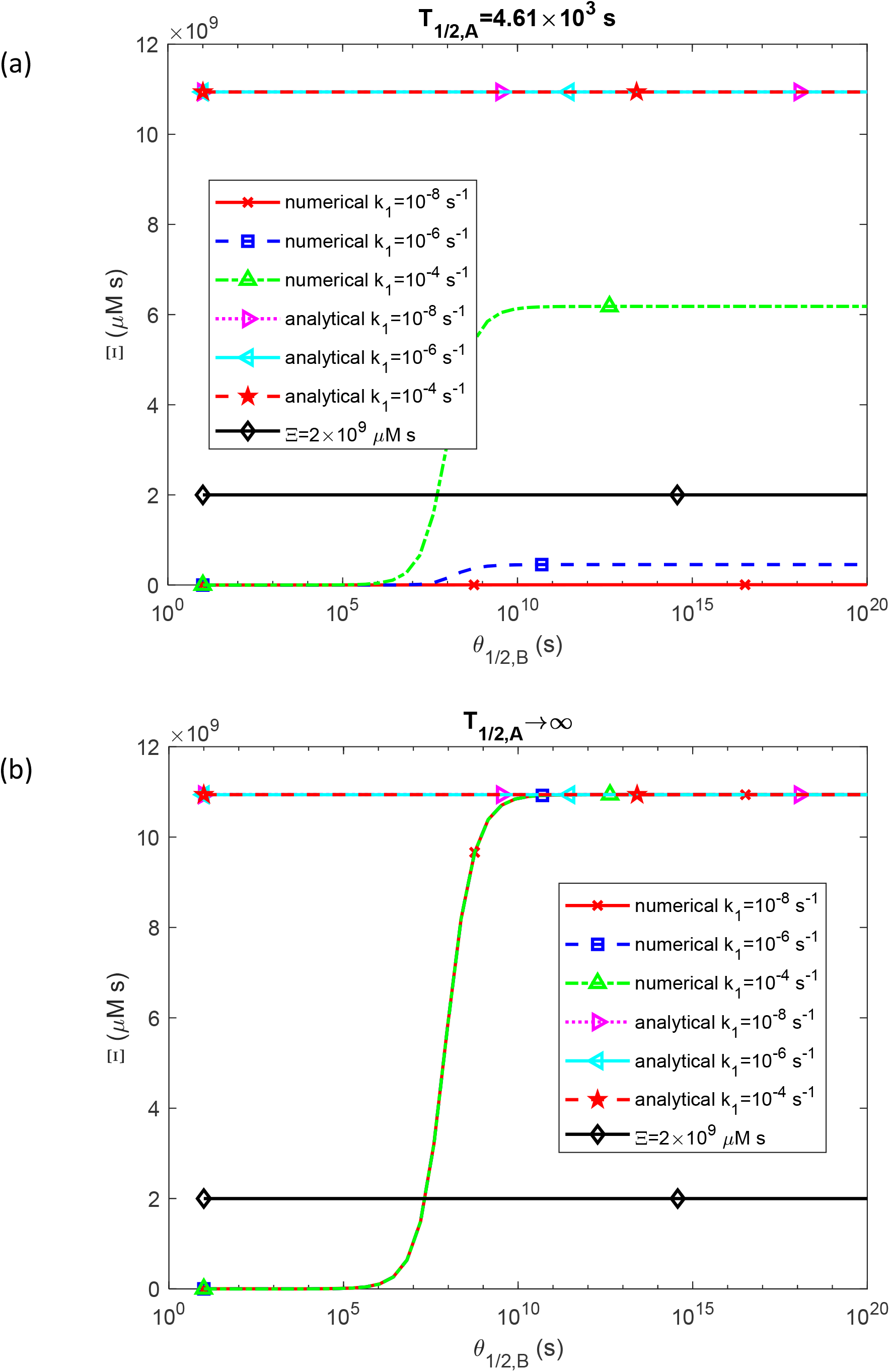
Accumulated toxicity of Aβ oligomers, Ξ, as a function of the half-deposition time of Aβ aggregates into senile plaques, *θ*_1/2,*B*_. Results are presented at *t* = *t* _*f*_ = 3.15 × 10^8^ s for three different values of *k*_1_. (a) *T*_1/ 2, *A*_ = 4.61×10^3^ s, (b) *T*_1/ 2, *A*_ →∞. Note that as *θ* _1/ 2,*B*_→ 0 all free aggregates immediately deposit into plaques, resulting in Ξ→0.

Figs. 7a and 7b, which show accumulated toxicity as a function of *θ*_1/2,*B*_ for various values of *k*_2_, tell a similar story. When *T*_1/ 2, *A*_ is finite (Fig. 7a), the Ξ= *f* (*θ*_1/2,*B*_) curves differ for different *k*_2_, but as *T*_1/2,*A*_ →∞ (Fig. 7b), the curves converge. As *θ*_1/2,*B*_ increases, some of these curves surpass the critical threshold of Ξ, 2 ×10^9^ µM·s, indicating that nearby neurons are likely to die.

**Fig. 7.**
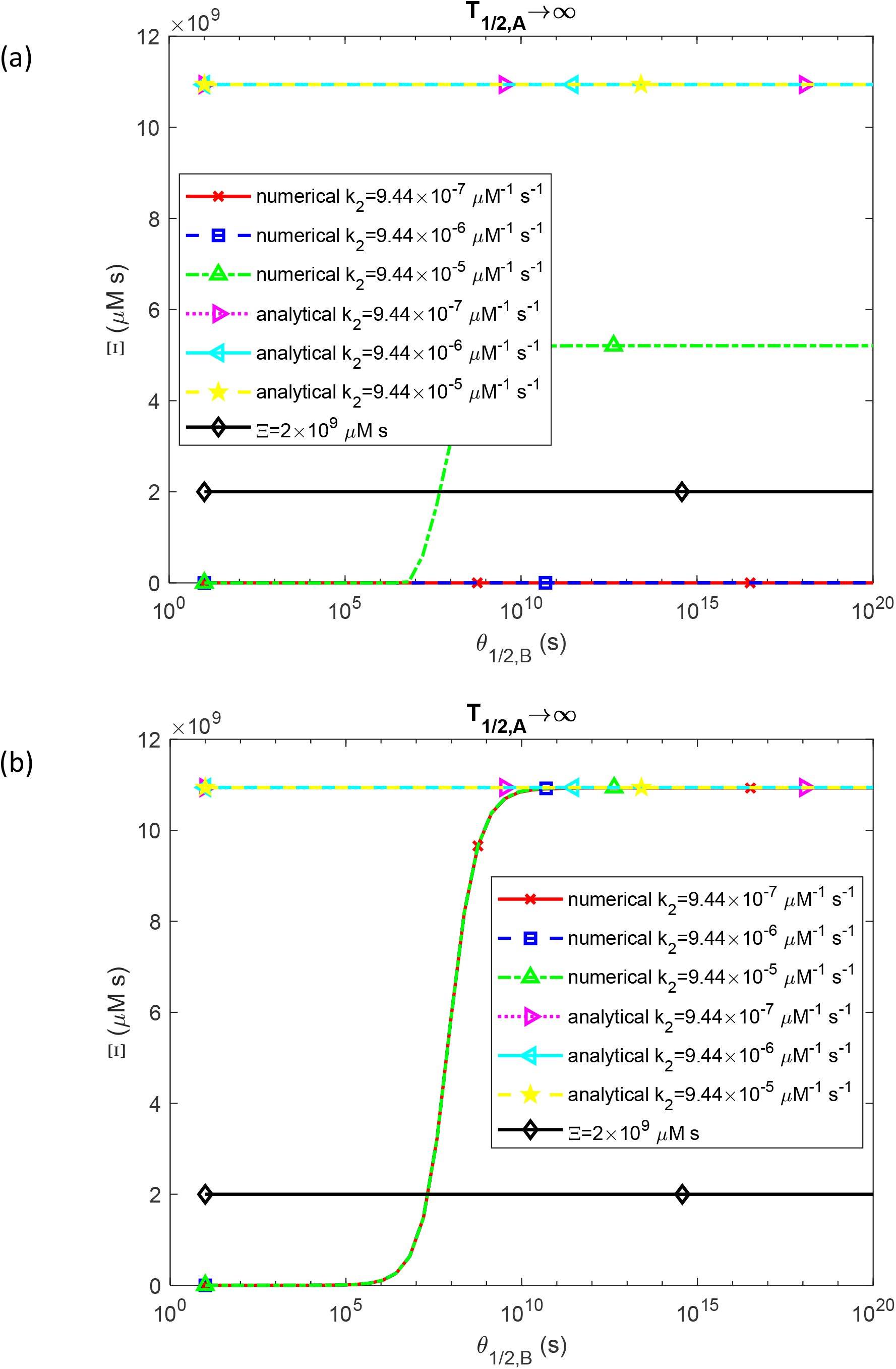
Accumulated toxicity of Aβ oligomers, Ξ, as a function of the half-deposition time of Aβ aggregates into senile plaques, *θ*_1/2,*B*_. Results are presented at *t* = *t* _*f*_ = 3.15 × 10^8^ s for three different values of *k*_2_. (a) *T*_1/ 2, *A*_ = 4.61×10^3^ s, (b) *T*_1/ 2, *A*_ →∞. Note that as *θ*_1/ 2,*B*_ → 0 all free aggregates immediately deposit into plaques, resulting in Ξ→0.

Figs. 8a and 8b illustrate how the curves for Ξ= *f* (*θ*_1/2,*B*_) vary with *q*_*A*,0_. Notably, the analytical solution given by Eq. (20) depends on *q*_*A*,0_, so each value of *q*_*A*,0_ corresponds to a different line in Figs. 8a and 8b. In Fig. 8b, which depicts the scenario where *T*_1/2,*A*_ →∞, each line for a different *q*_*A*,0_ approaches its own asymptotic limit, matching the value predicted by the analytical solution as *θ*_1/2,*B*_ →∞.

**Fig. 8.**
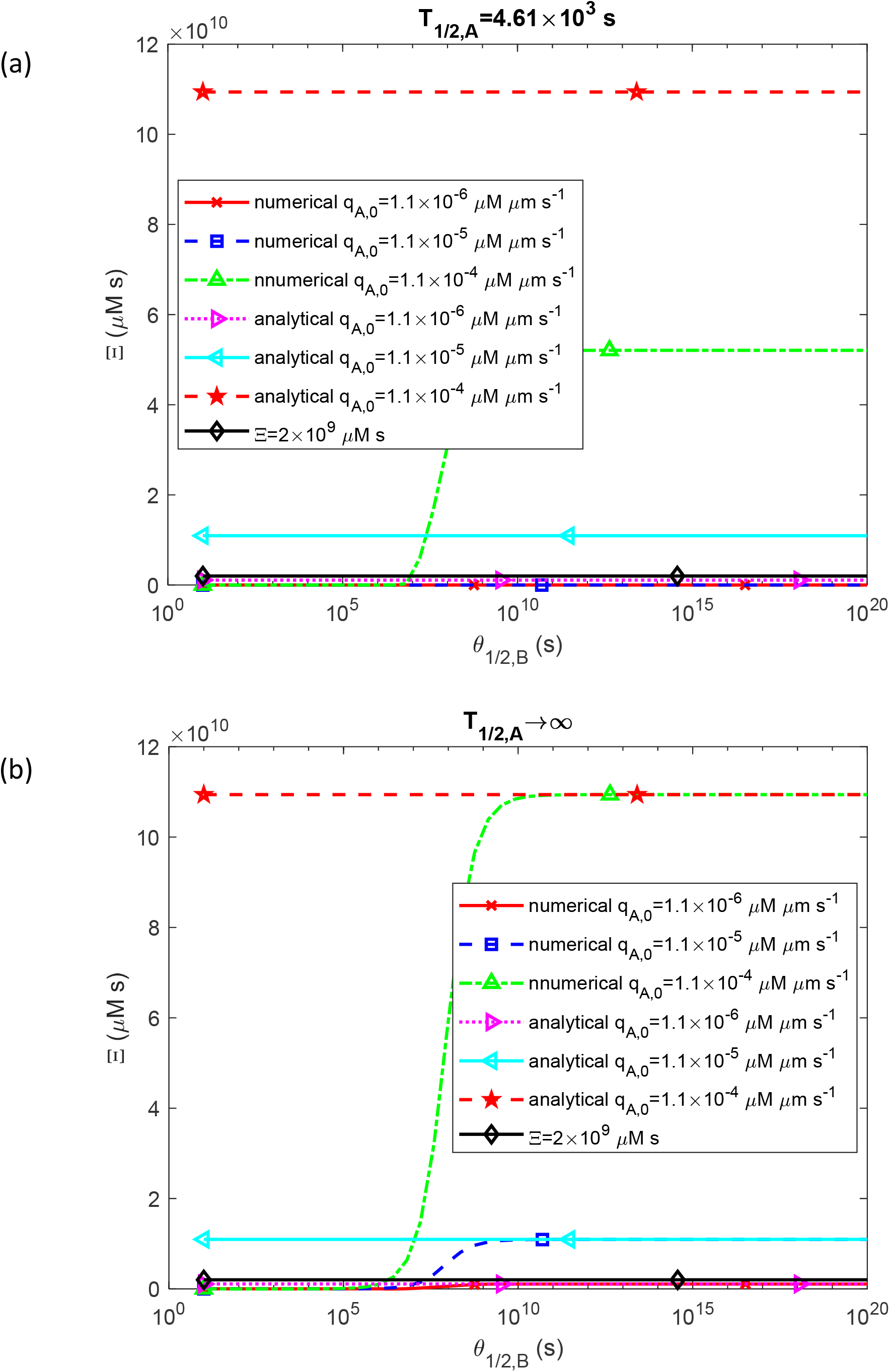
Accumulated toxicity of Aβ oligomers, Ξ, as a function of the half-deposition time of Aβ aggregates into senile plaques, *θ*_1/2,*B*_. Results are presented at *t* = *t* _*f*_ = 3.15 × 10^8^ s for three different values of *q*_*A*,0_. (a) *T*_1/ 2, *A*_ = 4.61×10^3^ s, (b) *T*_1/ 2, *A*_ →∞. Note that as *θ* _1/ 2,*B*_→ 0 all free aggregates immediately deposit into plaques, resulting in Ξ→0.

The sensitivity of accumulated toxicity, Ξ, to the nucleation rate constant, *k*, denoted as 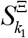, is positive (Fig. 9a). This indicates that increasing *k*_1_ results in a corresponding increase in Ξ. This aligns with the trend observed in Fig. 2a. From Fig. 9b, it is evident that in the scenario where *T*_1/ 2, *A*_ →∞ and *θ*_1/2,*B*_ →∞, 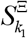 approaches the analytically predicted value of zero over time (see Eq. (33)).

**Fig. 9.**
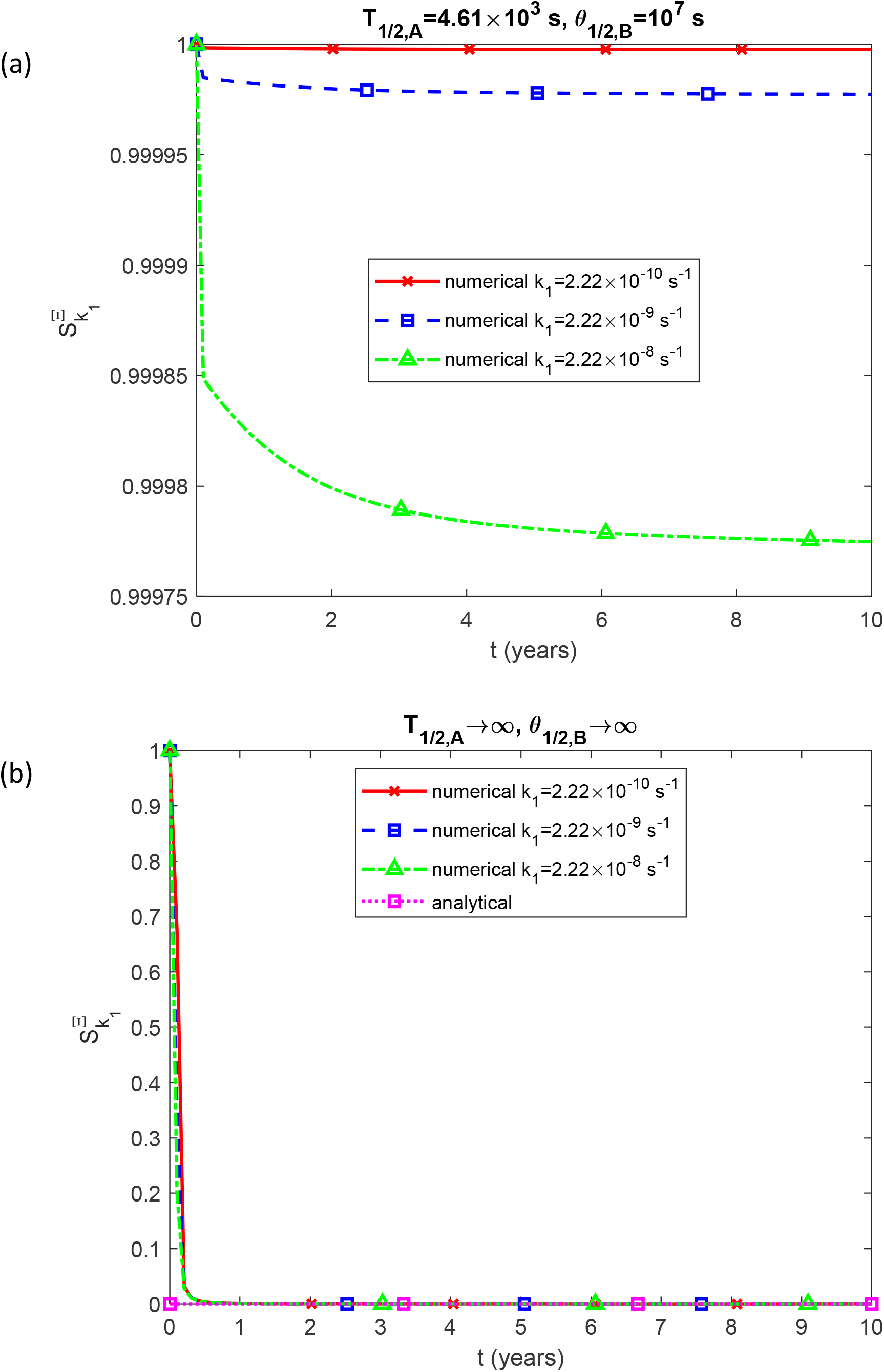
Sensitivity of the accumulated toxicity, Ξ, to the rate constant that describes the nucleation of Aβ aggregates, *k*_1_. Results are shown for three different values of *k*_1_. (a) *T*_1/ 2, *A*_ = 4.61×10^3^ s and *θ*_1/2,*B*_ =10^7^ s, (b) *T*_1/ 2, *A*_ →∞ and *θ*_1/2,*B*_ →∞.

The accumulated toxicity, Ξ, exhibits a positive sensitivity to the rate constant describing autocatalytic growth, *k*_2_ (Fig. 10a), so a greater *k*_2_ leads to an increase in Ξ. This trend is consistent with the result depicted in Fig. 3a. As shown in Fig. 10b, as *T*_1/ 2, *A*_ →∞ and, 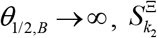 converges to the zero value predicted by the analytical solution as time progresses (see Eq. (34)).

**Fig. 10.**
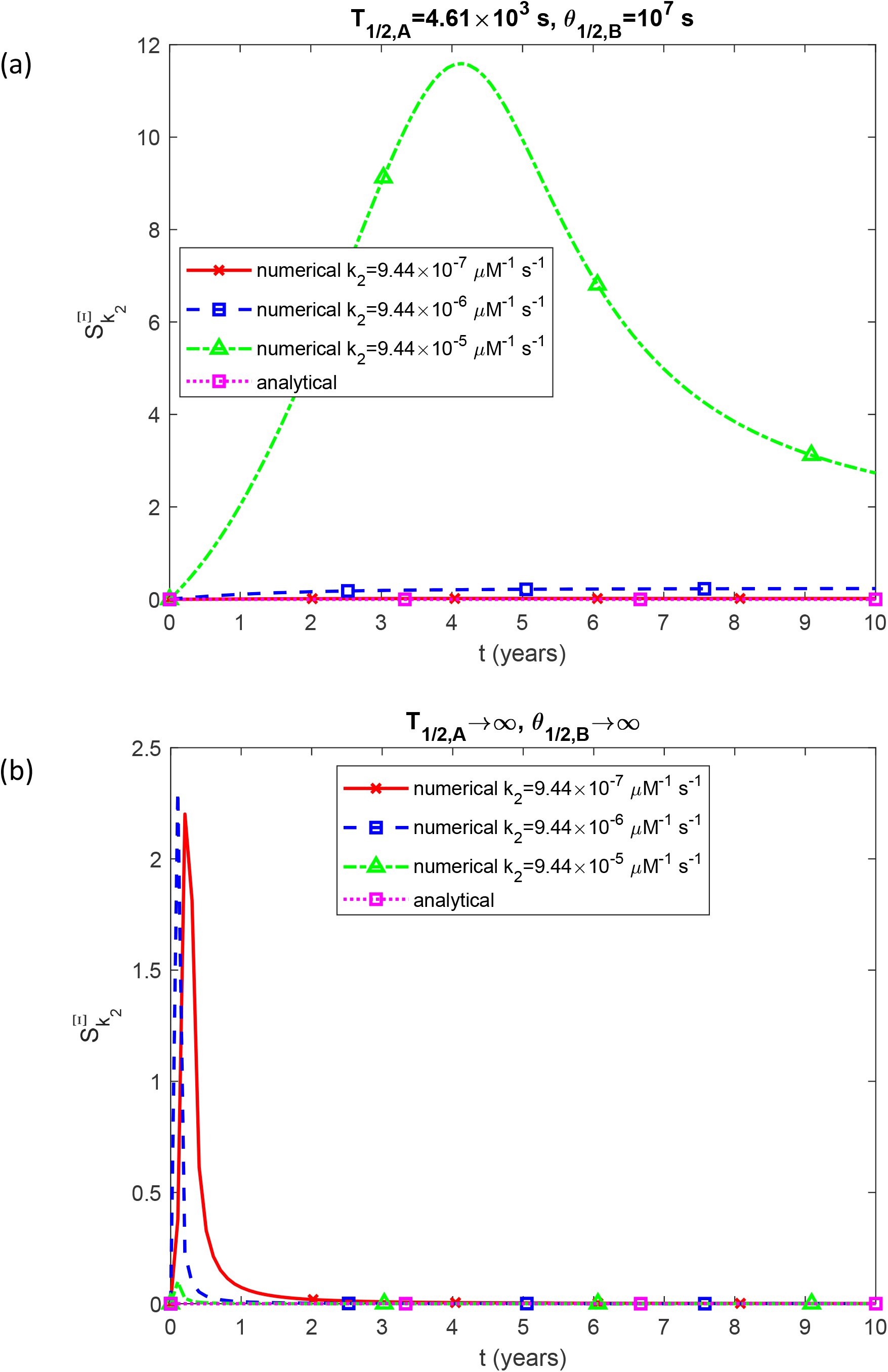
Sensitivity of the accumulated toxicity, Ξ, to the rate constant that describes the autocatalytic growth Aβ aggregates, *k*_2_. Results are shown for three different values of *k*_2_. (a) *T*_1/ 2, *A*_ = 4.61×10^3^ s and *θ* _1/2,*B*_=10^7^ s, (b) *T*_1/ 2, *A*_ →∞ and *θ* _1/2,*B*_→∞.

The sensitivity of accumulated toxicity, Ξ, to the flux of Aβ monomers, *q*_*A*,0_, is also positive (Fig. 11a). Hence, increasing *q*_*A*,0_ leads to a higher Ξ. This result aligns with the observation in Fig. 4a. Fig. 11b shows that as *T*_1/ 2, *A*_ →∞ and, 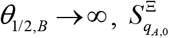 reaches the analytically predicted value of unity as time increases (see Eq. (35)).

**Fig. 11.**
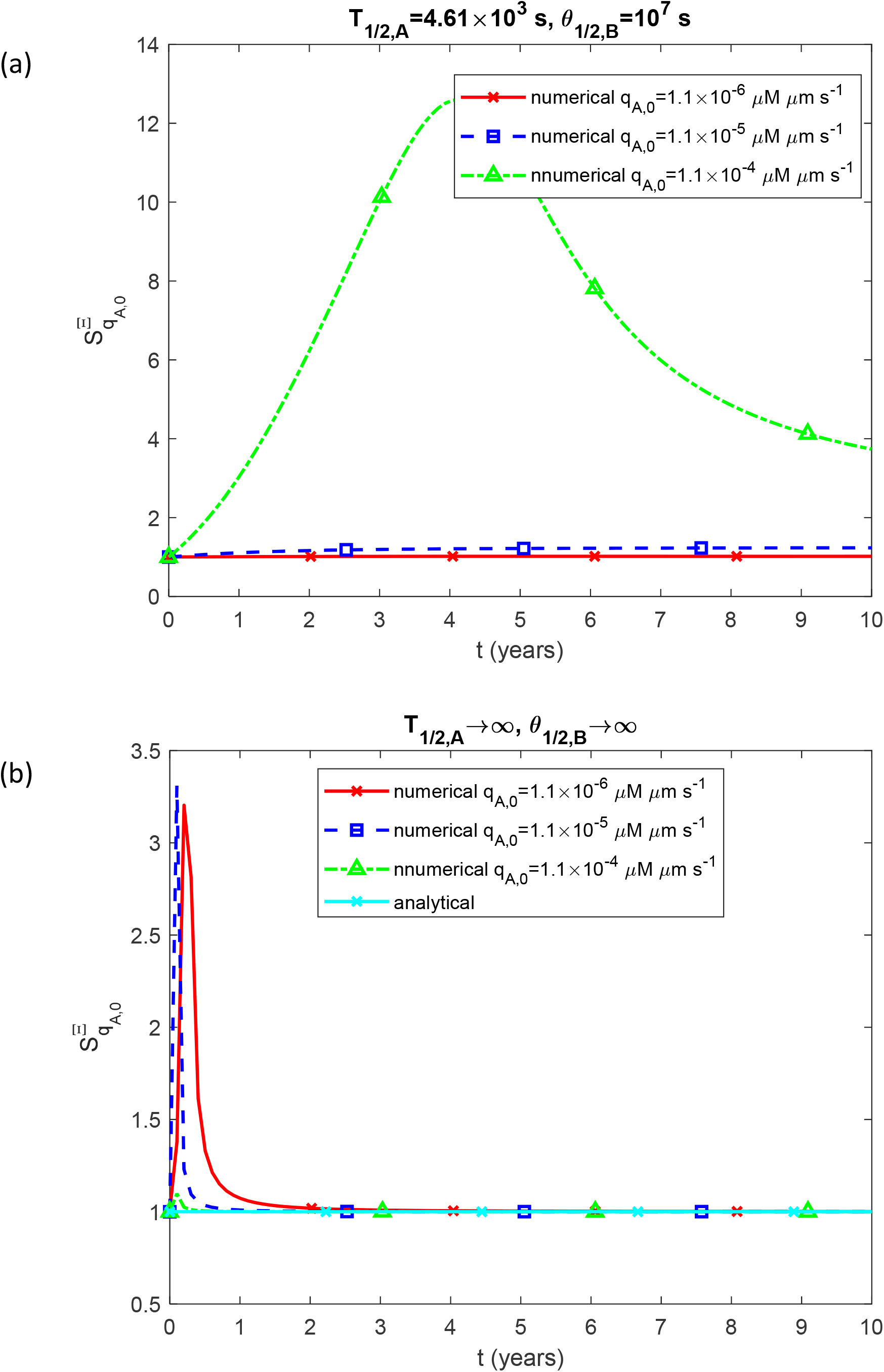
Sensitivity of the accumulated toxicity, Ξ, to the flux of Aβ monomers, *q*_*A*,0_. Results are shown for three different values of *q*_*A*,0_. (a) *T*_1/ 2, *A*_ = 4.61×10^3^ s and *θ*_1/2,*B*_ =10^7^ s, (b) *T*_1/ 2, *A*_ →∞ and *θ*_1/2,*B*_→∞.

## 6. Discussion, limitations of the model, and future directions

A criterion for characterizing the accumulated toxicity of Aβ oligomers at the microscale is developed, along with a proposed critical threshold beyond which nearby neurons are expected to die. This criterion allows to quantify the Aβ oligomer toxicity at the cellular scale. Establishing such a criterion is crucial for advancing AD research, as most current studies investigating neuronal damage at the cellular scale primarily focus on qualitative descriptions of the neuronal damage caused by Aβ aggregation, with limited attention to quantitative analysis.

If Aβ monomers and aggregates have an infinite half-life, then accumulated toxicity, Ξ, shows an S-shaped growth pattern as the half-deposition time of Aβ aggregates into senile plaques, *θ*_1/2,*B*_, increases. Once *θ*_1/2,*B*_ becomes sufficiently large, Ξ levels off and aligns with the value predicted by the analytical solution for Ξ.

The analytical solution in Eq. (20), which is valid for a small rate of deposition of free aggregates into senile plaques (large *θ*_1/2,*B*_) and an infinitely long half-life of Aβ monomers and aggregates, indicates that accumulated toxicity is proportional to the square of the duration of the senile plaque formation process.

This implies that, given enough time (assuming a long enough lifespan), the accumulated toxicity will eventually surpass the critical threshold, leading to neuron death. However, the analytical solution is only valid in the scenario of infinitely long half-life of Aβ monomers and free aggregates, indicating a malfunctioning degradation machinery for both monomers and aggregates. Therefore, the model suggests that if the protein degradation machinery is impaired, AD becomes inevitable—it is only a matter of time before neurons begin to die. The only way to prevent this is by ensuring that the degradation machinery for Aβ remains functional. This aligns with a hypothesis proposed by several researchers, suggesting that an imbalance between Aβ production and clearance can lead to accumulation of misfolded Aβ in the brain [40].

The quadratic dependence on the duration of the process predicted by the analytical solution (Eq. (20)) suggests that the toxicity accumulation starts off slowly but accelerates over time. This may explain why AD symptoms can take years to manifest.

Accumulated toxicity, Ξ, exhibits positive sensitivity to the nucleation rate constant, *k*_1_, the autocatalytic growth rate constant, *k*_2_, and the flux of Aβ monomers, *q*_*A*,0_. For each of these cases, the sensitivity converges to the analytically predicted values when the half-deposition time of Aβ aggregates into senile plaques, *θ*_1/2,*B*_, is large, and when the half-lives for Aβ monomers and aggregates, *T*_1/2, *A*_ and *T*_1/2,*B*_, respectively, are infinitely long.

The toxicity of Aβ plaques might be attributed to the high concentration of oligomers accumulating near the plaques due to their deposition. To test this hypothesis, it would be beneficial to develop a model that quantifies oligomer concentration in the diffusion boundary layer near a plaque’s surface. This model could also assess the likelihood that these elevated oligomer levels could trigger inflammatory responses, resulting in neuronal damage. Additionally, developing a model to simulate the interactions between a diverse array of oligomers and cellular membranes would be valuable.

The F-W model lumps all aggregates, such as dimers and trimers, into a single phase representing all free amyloid-converted peptides (phase *B* in Eqs. (1) and (2)) with a concentration denoted as *C*_*B*_. As a result, it does not differentiate between various types of Aβ oligomers. Future research should address this limitation by using more detailed models that can simulate the formation of different Aβ oligomers.

It would be also important to establish a correlation between the criterion developed in this paper and the severity of cognitive decline in patients. A significant next step in this research would be to develop a criterion that accounts for the combined neural damage caused by Aβ oligomers and pathological tau.

## Abbreviations

Aβ: amyloid beta
AD: Alzheimer’s disease
CV: control volume
F-W: Finke-Watzky

## Data accessibility

The source MATLAB code used to generate the figures in this paper is publicly available at Github: https://github.com/ikuznet1/Kuznetsov_Alzheimer_2025/tree/main. Any additional information required to reanalyze the data reported in this paper is available from the authors upon reasonable request. Additional data are also provided in electronic supplementary material.

## Authors’ contributions

AVK is the sole author of this paper. Competing interests. The author declares no competing interests.

## Supplemental Materials

## S1. Dimensionless equations

### S1.1. Dimensionless form of Eqs. (4)-(8)

The model described by Eqs. (4)-(7) is reformulated in dimensionless form as shown below, where asterisks denote dimensionless quantities:

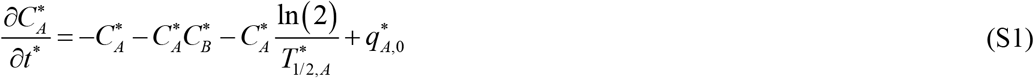

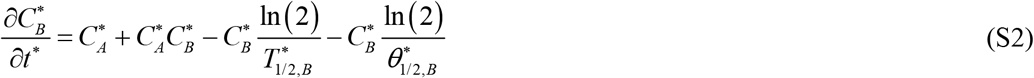

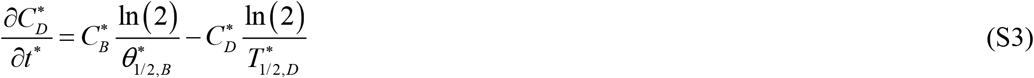

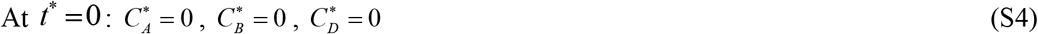

The independent dimensionless variable in the model is given in Table S1, with the dependent dimensionless variables summarized in Table S2. Table S3 provides a summary of the dimensionless parameters used in the model.

The parameter representing the accumulated toxicity of Aβ, as given by Eq. (8), can be reformulated in terms of dimensions quantities as follows:

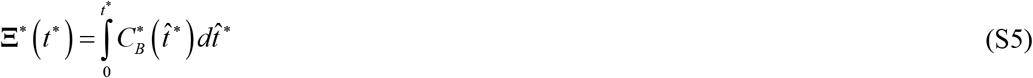

Where

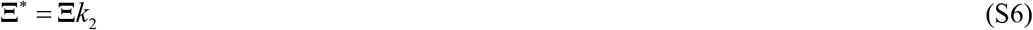

### S1.2. Analytical solution of Eqs. (15) and (16)

Combining Eqs. (15) and (16) yields

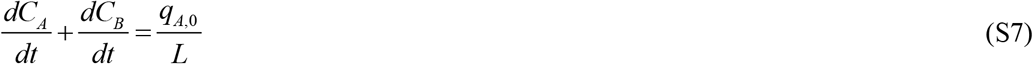

By integrating Eq. (S7) over time and applying the initial conditions specified in Eq. (7a,b), the following result is derived:

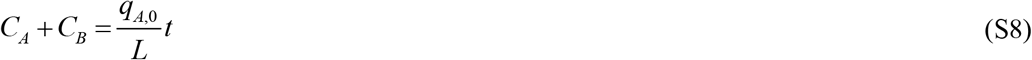

The increase in *C*_*A*_ + *C*_*B*_ over time is due to the influx of Aβ monomers through the left boundary of the CV. By substituting Eq. (S8) into Eq. (16) to eliminate *C*_*A*_, the following expression is obtained:

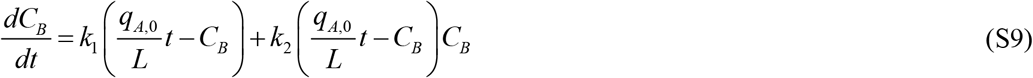

Eq. (S9) is similar to the equation studied in ref. [31]. To obtain the exact solution of Eq. (S9) with the initial condition specified in Eq. (7b), Mathematica 13.3 (Wolfram Research, Champaign, IL) was utilized, employing the DSolve function followed by the FullSimplify function. The derived solution is as follows:

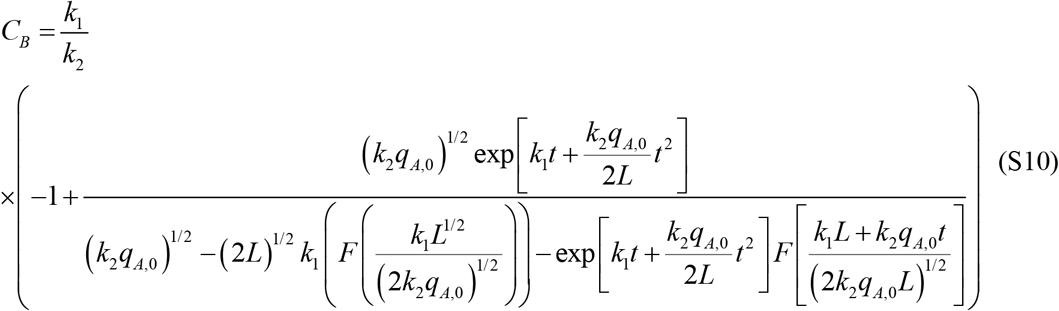

In this equation, *F* (*x*) represents Dawson’s integral:

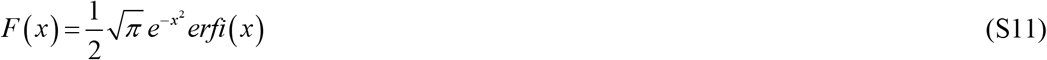

where *erfi* (*x*) is the imaginary error function.

As *t* → 0, Eq. (S10) shows that 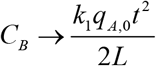, suggesting that at early times, the concentration of Aβ aggregates is linearly dependent on the nucleation rate constant, *k*_1_. Using Eq. (S8), the expression for *C*_*A*_ can be derived as:

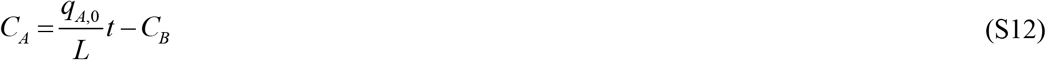

To estimate the maximum possible plaque radius, one can assume that all aggregates deposit into plaques. However, this assumption conflicts with the condition *θ*_1/2,*B*_ →∞, so the maximum radius would be reached at some finite value of *θ*_1/ 2, *B*_ (but not at zero).

If the approximate solution given by Eqs. (18) and (19) is used, the following result is obtained.

Renaming *C*_*B*_ given in Eq. (19) and substituting it into Eq. (14) yield

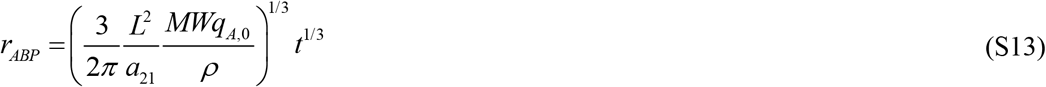

which represents the “cube root of time” hypothesis, originally formulated in ref. [14].

### S1.3. Numerical solution

Eqs. (4)–(6), along with their initial conditions in Eq. (7), were numerically solved using MATLAB’s ODE45 solver (MATLAB R2020b, MathWorks, Natick, MA, USA), which is widely recognized for its accuracy and robustness. The relative and absolute tolerances, RelTol and AbsTol, were set to 1e-10 to ensure high precision in the solutions.

## S2. Supplementary tables

**Table S1.**
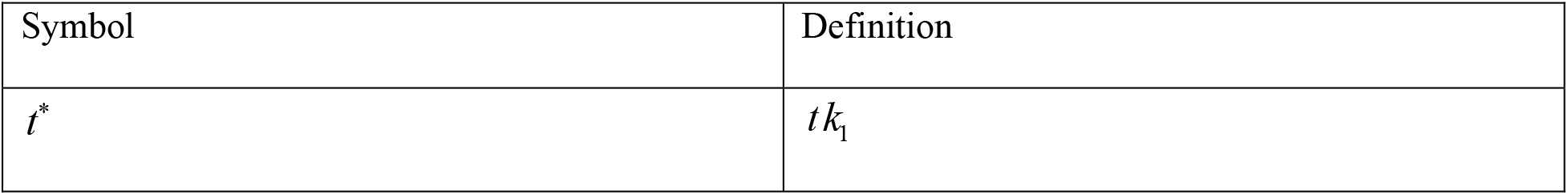
Dimensionless independent variables used in the model.

| Symbol | Definition |
| --- | --- |
| $t^*$ | $t k_1$ |

**Table S2.** Dimensionless dependent variables used in the model.

| Symbol | Definition |
| --- | --- |
| $C_A^*$ | $C_A k_2 / k_1$ |
| $C_B^*$ | $C_B k_2 / k_1$ |
| $C_D^*$ | $C_D k_2 / k_1$ |

**Table S3.** Dimensionless parameters used in the model.

| Symbol | Definition |
| --- | --- |
| $q_{A,0}^*$ | $q_{A,0} k_2 / (k_1^2 L)$ |
| $t_f^*$ | $t_f k_1$ |
| $T_{1/2,A}^*$ | $T_{1/2,A} k_1$ |
| $T_{1/2,B}^*$ | $T_{1/2,B} k_1$ |
| $T_{1/2,D}^*$ | $T_{1/2,D} k_1$ |
| $\theta_{1/2,B}^*$ | $\theta_{1/2,B} k_1$ |
| $\rho^*$ | $\frac{\rho}{MW} a_{21} \frac{k_2}{k_1}$ |

## S3. Supplementary figures

**Fig. S1.**
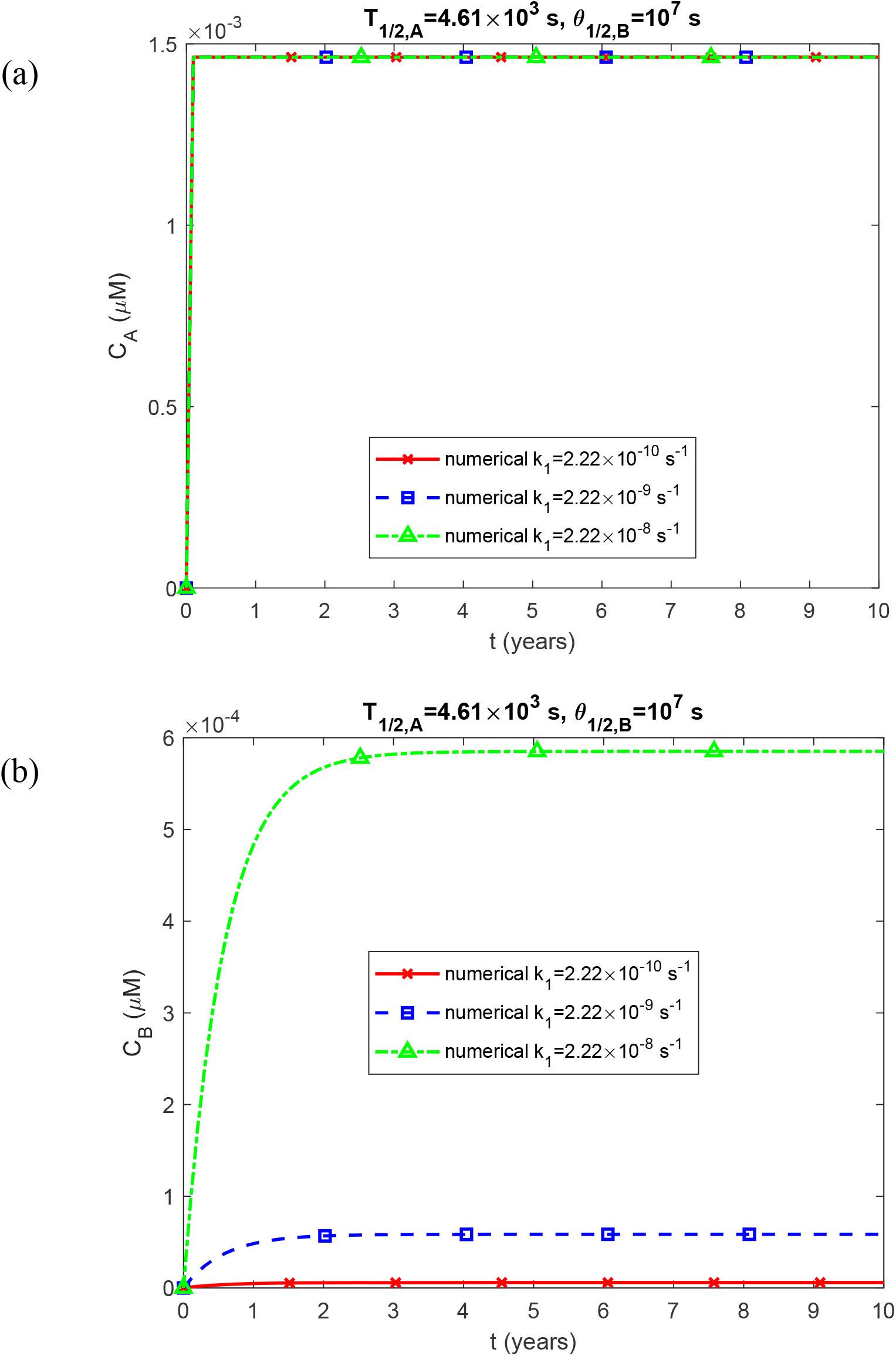
Molar concentration of Aβ monomers, *C*_*A*_ (a); and free Aβ aggregates, *C*_*B*_ (b) as a function of time (in years). Results are shown for three different values of *k*_1_. The scenario with a finite half-life of Aβ monomers (_1/ 2, *A*_*T* = 4.61×10^3^ s) and an intermediate value of half-deposition time of Aβ aggregates into senile plaques (*θ*_1/2,*B*_ =10^7^ s).

**Fig. S2.**
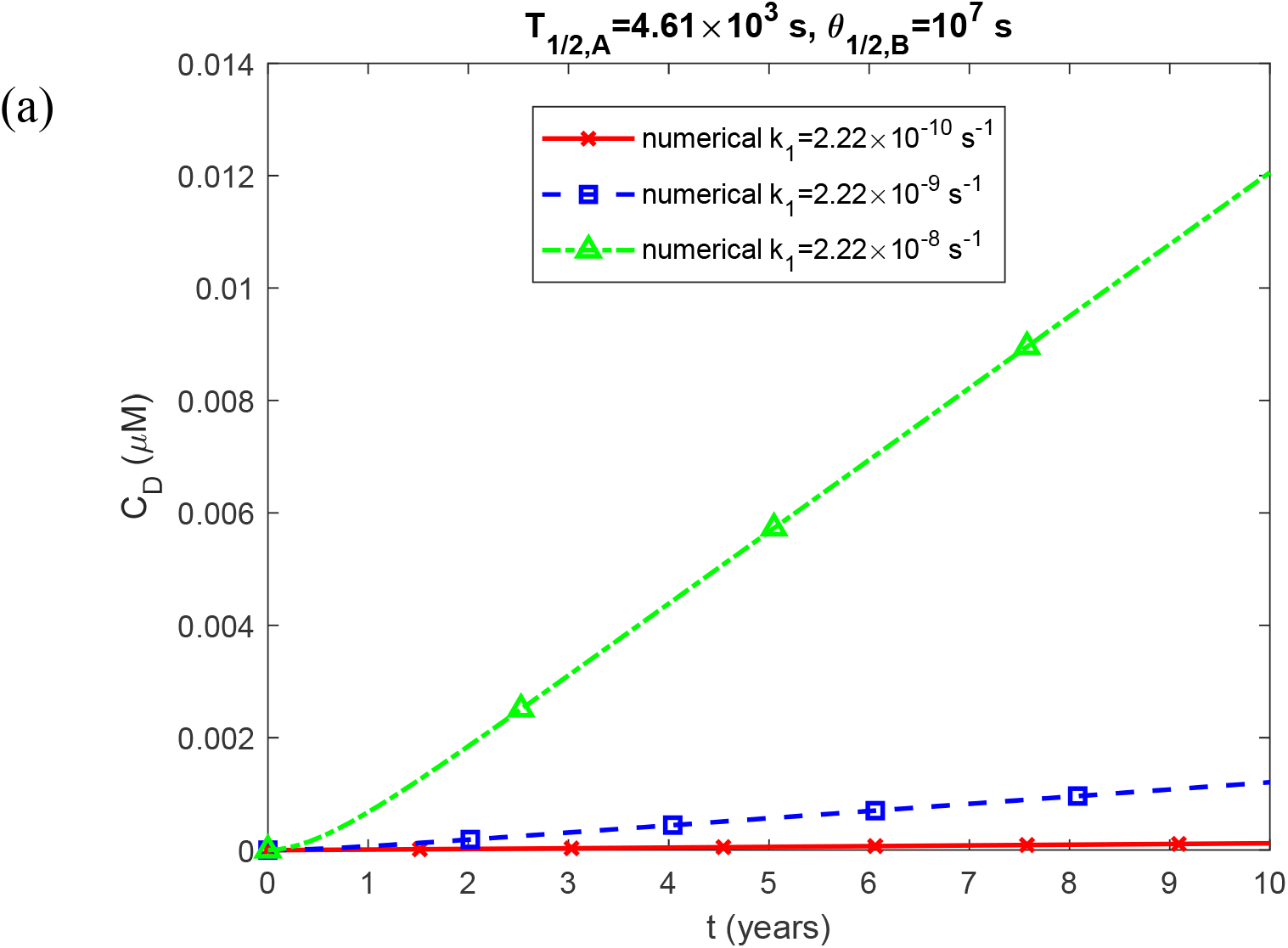
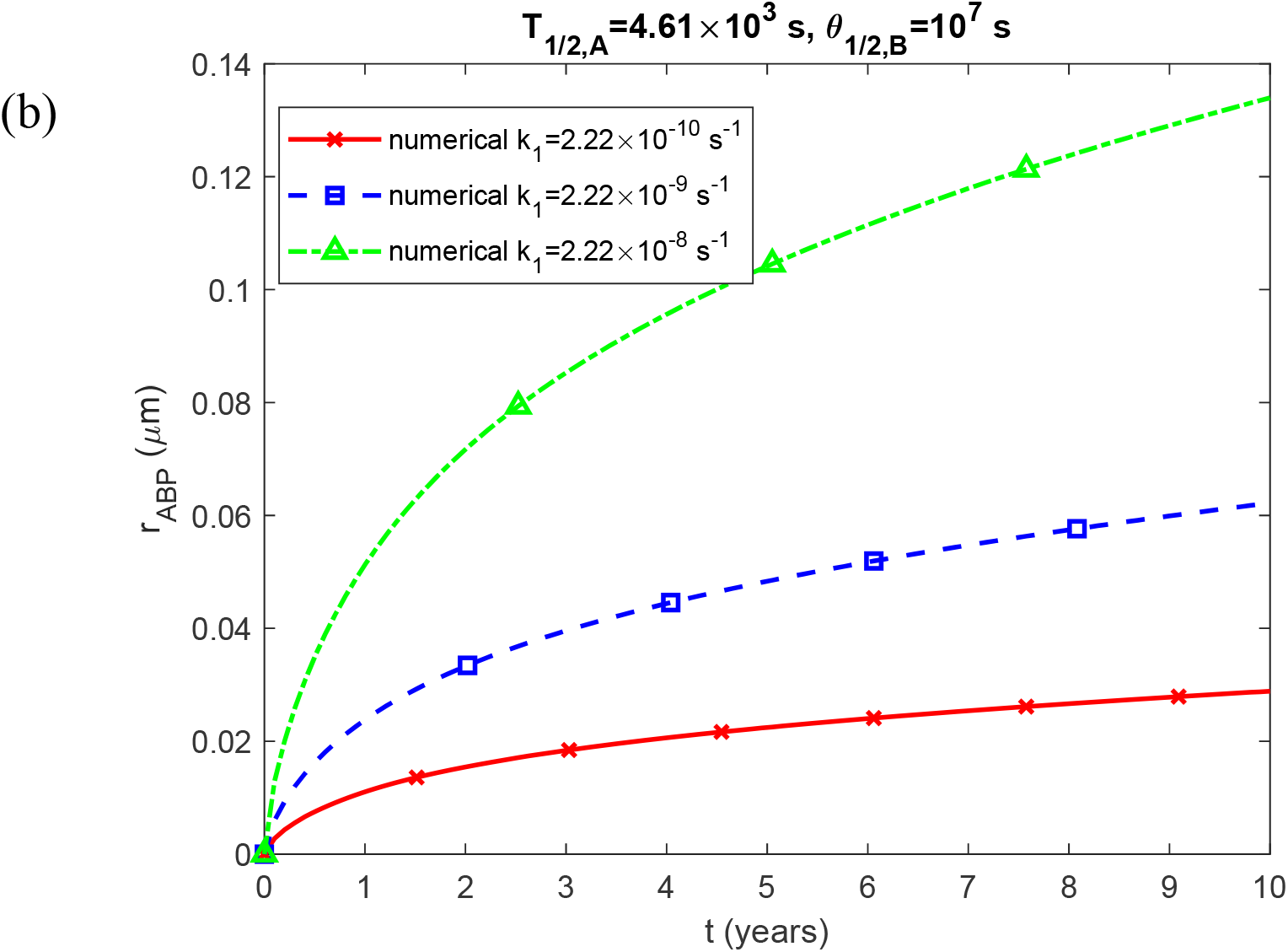
Molar concentration of Aβ aggregates deposited in plaques, *C*_*D*_ (a); and the radius of a growing Aβ plaque, *r*_*ABP*_ (b); as a function of time (in years). Results are shown for three different values of *k*_1_. The scenario with a finite half-life of Aβ monomers (*T*_1/ 2, *A*_ = 4.61×10^3^ s) and an intermediate value of half-deposition time of Aβ aggregates into senile plaques (*θ* _1/2,*B*_=10^7^ s).

**Fig. S3.**
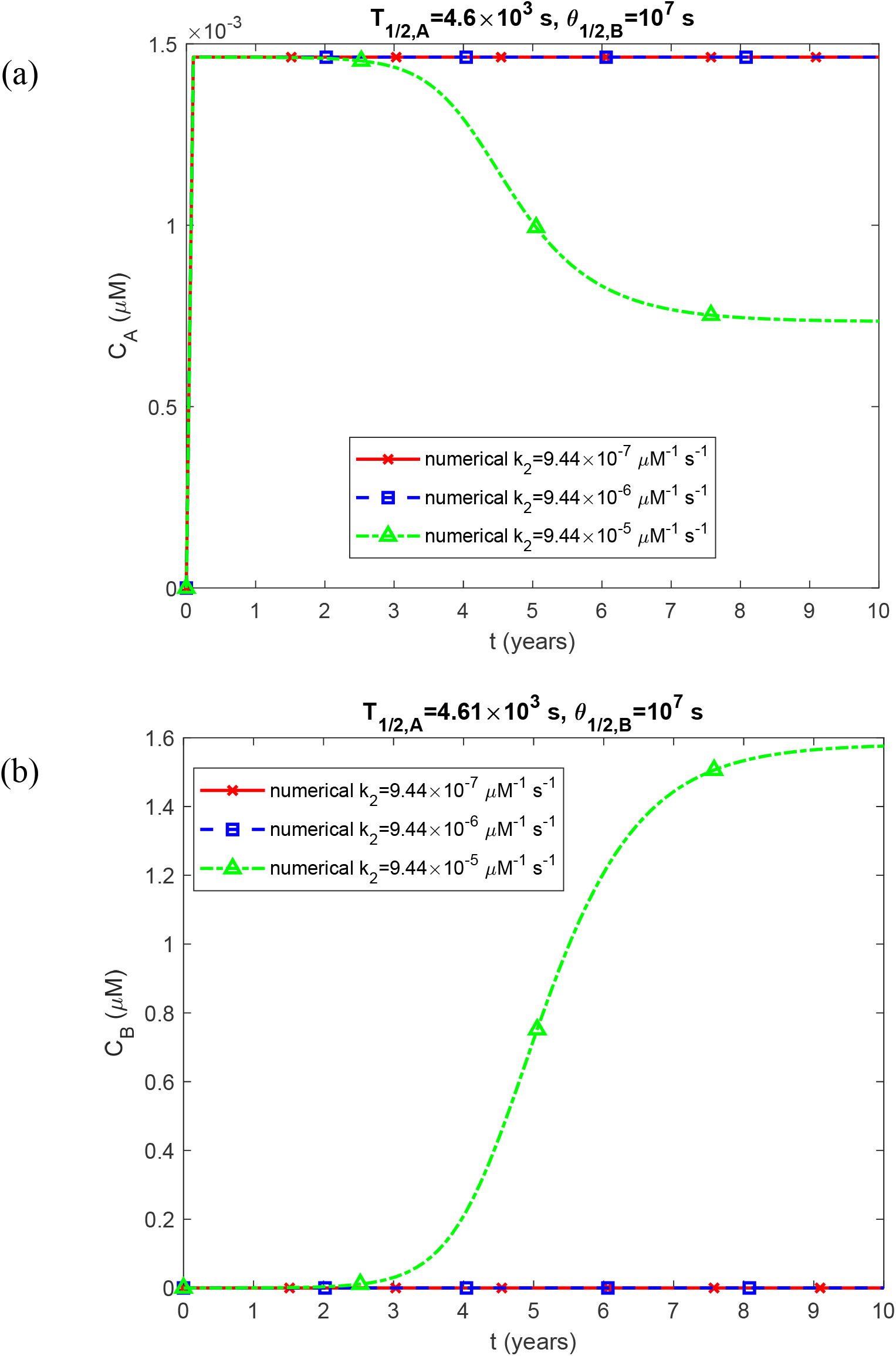
Molar concentration of Aβ monomers, *C*_*A*_ (a); and free Aβ aggregates, *C*_*B*_ (b) as a function of time (in years). Results are shown for three different values of *k*_2_. The scenario with a finite half-life of Aβ monomers (*T*_1/ 2, *A*_ = 4.61×10^3^ s) and an intermediate value of half-deposition time of Aβ aggregates into senile plaques (*θ* _1/2,*B*_=10^7^ s).

**Fig. S4.**
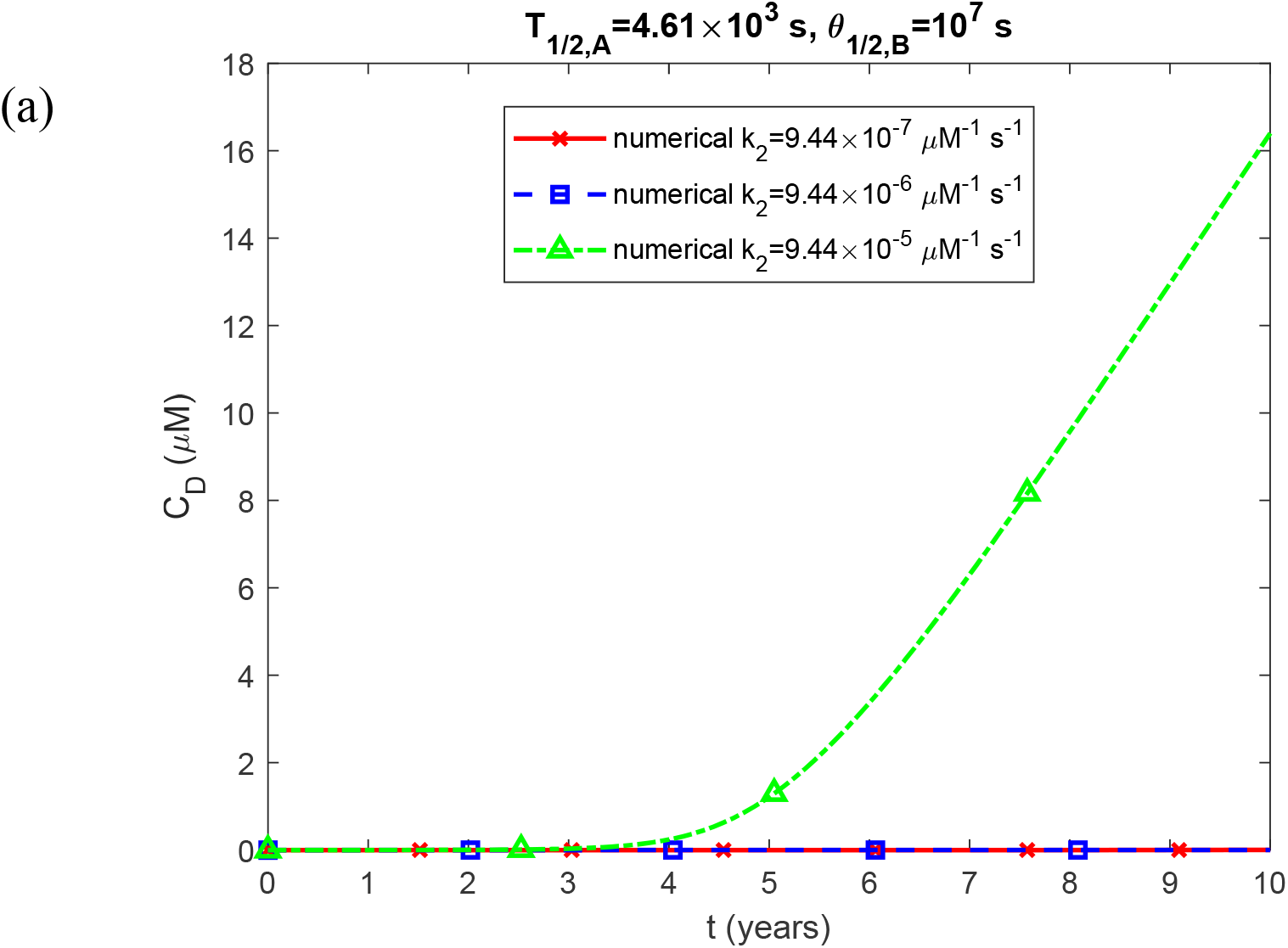
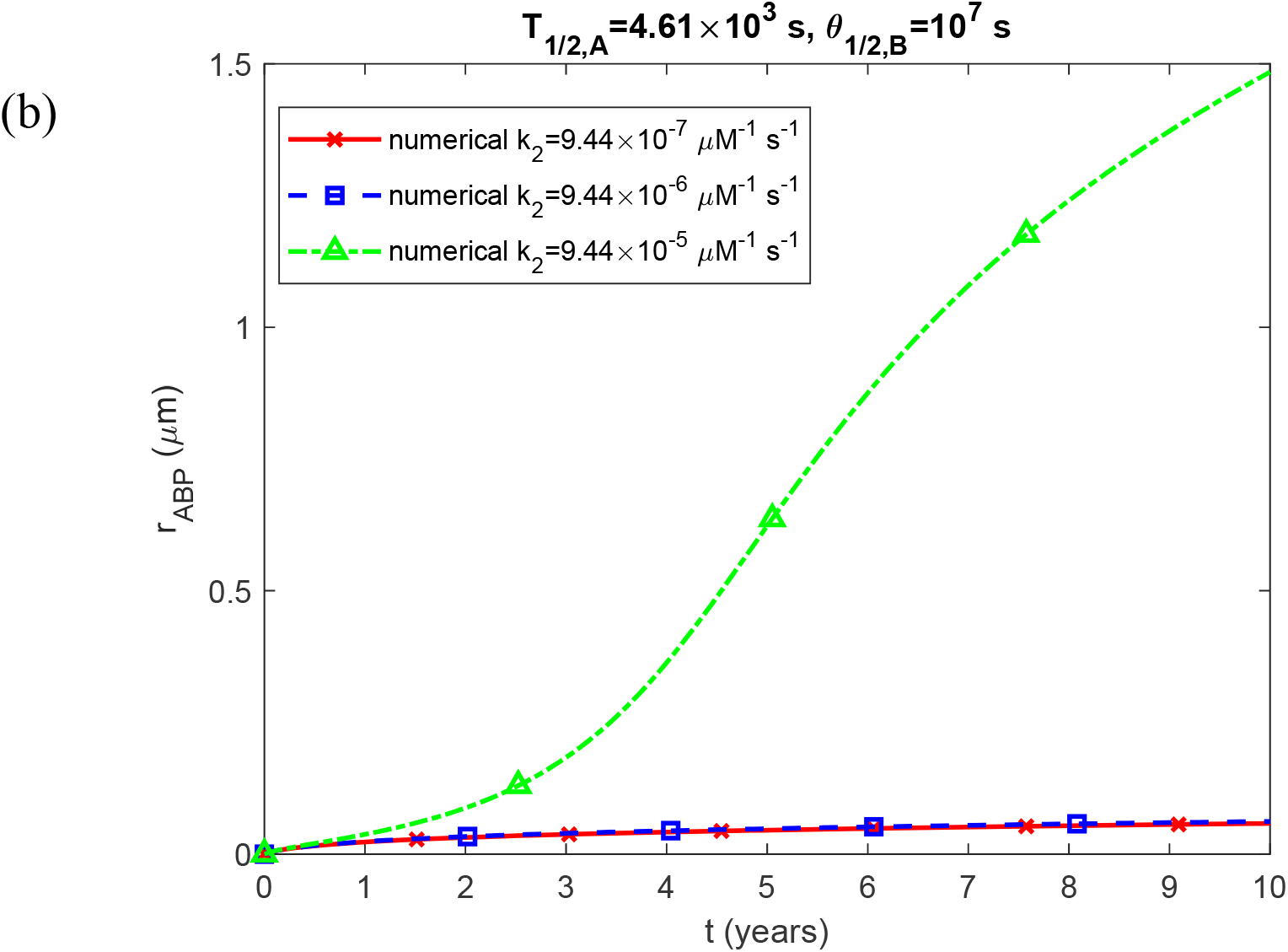
Molar concentration of Aβ aggregates deposited in plaques, *C*_*D*_ (a); and the radius of a growing Aβ plaque, *r*_*ABP*_ (b); as a function of time (in years). Results are shown for three different values of *k*_2_. The scenario with a finite half-life of Aβ monomers (*T*_1/ 2, *A*_ = 4.61×10^3^ s) and an intermediate value of half-deposition time of Aβ aggregates into senile plaques (*θ* _1/2,*B*_=10^7^ s).

**Fig. S5.**
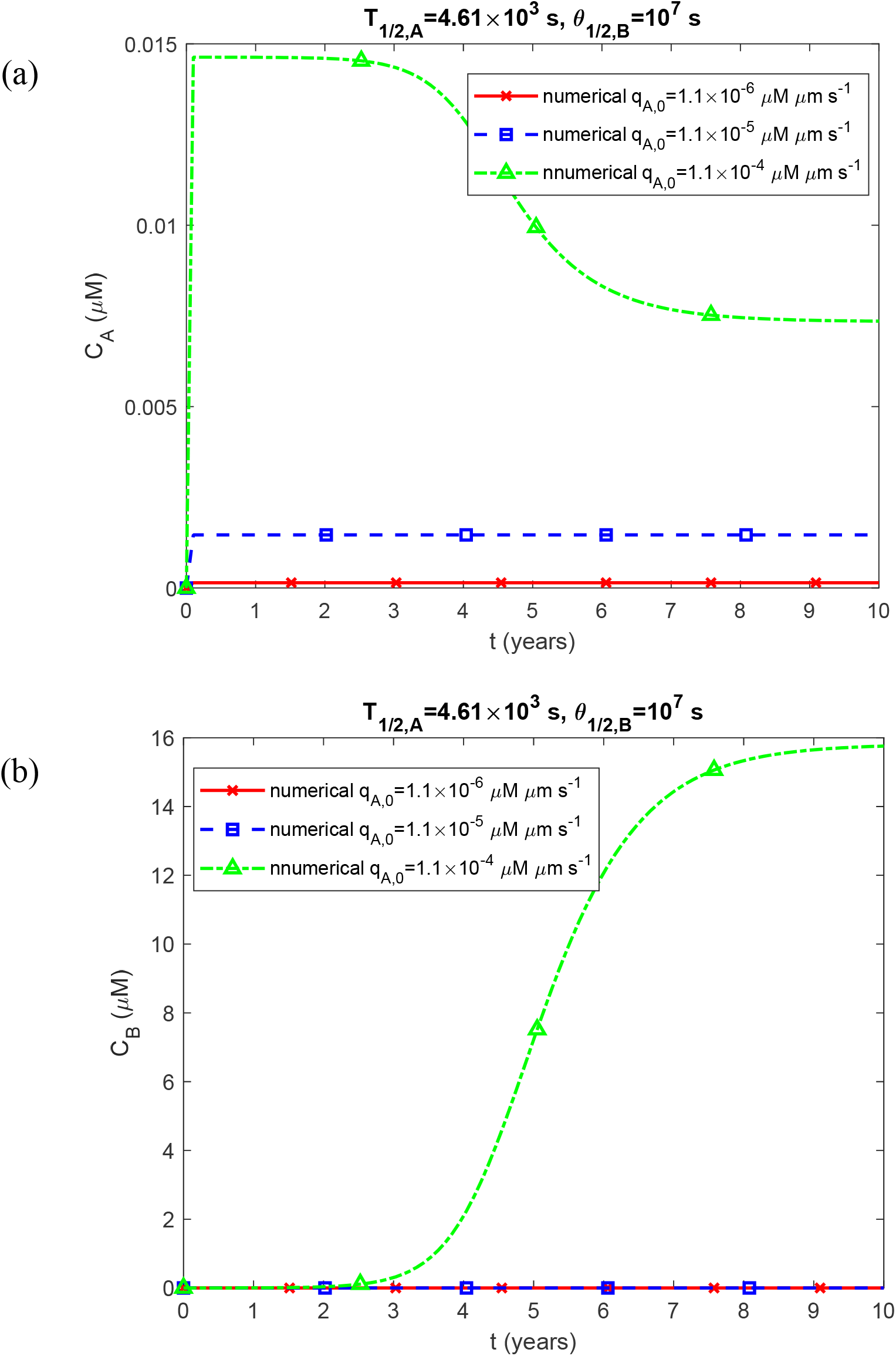
Molar concentration of Aβ monomers, *C*_*A*_ (a); and free Aβ aggregates, *C*_*B*_ (b) as a function of time (in years). Results are shown for three different values of *q*_*A*,0_. The scenario with a finite half-life of Aβ monomers (*T*_1/ 2, *A*_ = 4.61×10^3^ s) and an intermediate value of half-deposition time of Aβ aggregates into senile plaques (*θ*_1/2,*B*_ =10^7^ s).

**Fig. S6.**
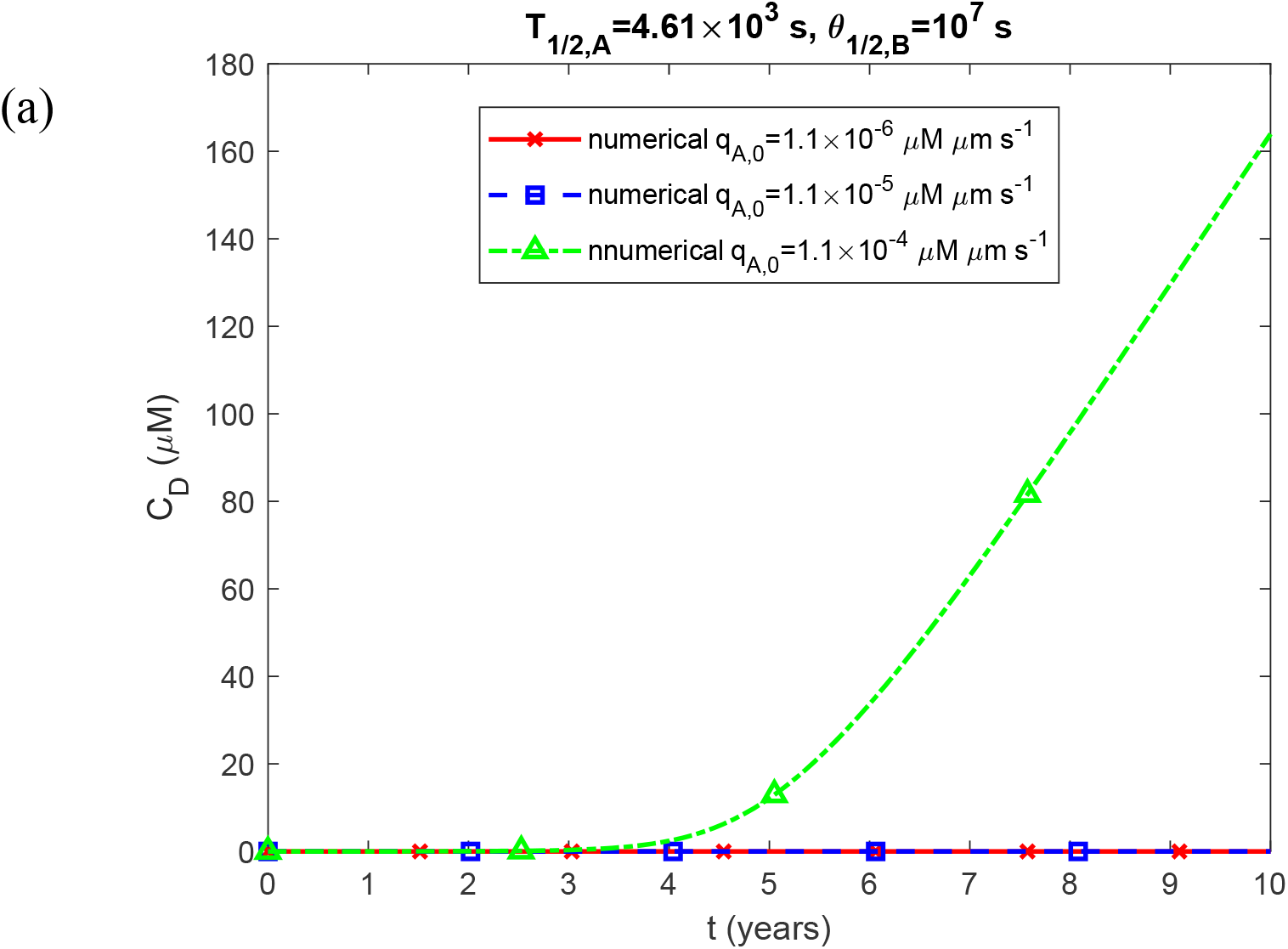
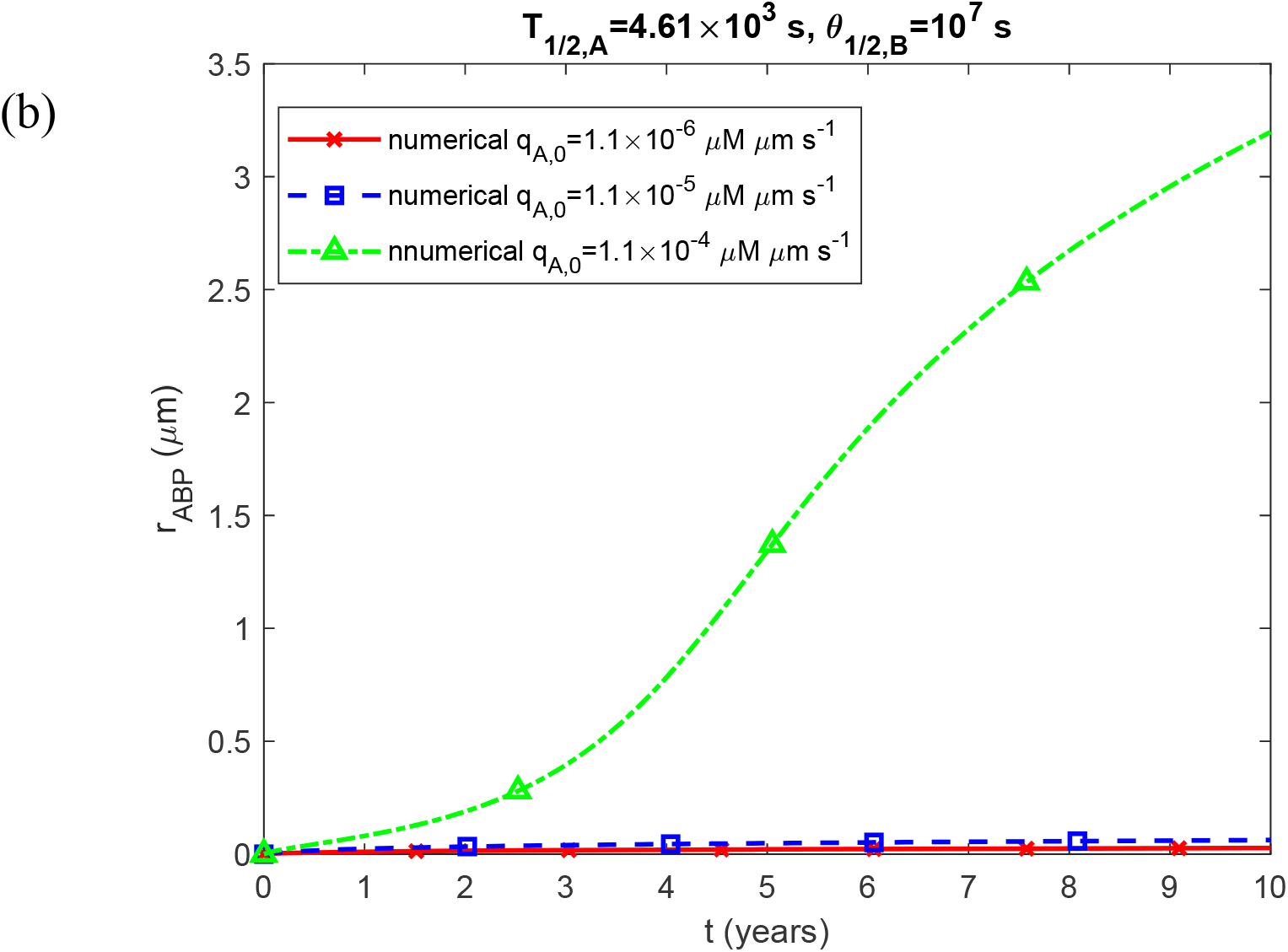
Molar concentration of Aβ aggregates deposited in plaques, *C*_*D*_ (a); and the radius of a growing Aβ plaque, *r*_*ABP*_ (b); as a function of time (in years). Results are shown for three different values of *q*_*A*,0_. The scenario with a finite half-life of Aβ monomers (*T*_1/ 2, *A*_ = 4.61×10^3^ s) and an intermediate value of half-deposition time of Aβ aggregates into senile plaques (*θ*_1/2,*B*_ =10^7^ s).

